# Evolution of novel mimicry polymorphisms through Haldane’s sieve and rare recombination

**DOI:** 10.1101/2024.07.24.605018

**Authors:** Riddhi Deshmukh, Saurav Baral, Athulya Girish Kizhakke, Muktai Kuwalekar, Krushnamegh Kunte

## Abstract

Origins of phenotypic novelty represent a paradox. Maintenance of distinct, canalized morphs usually requires a complex array of polymorphisms, whose co-retention requires a genetic architecture resistant to recombination, involving inversions and master regulators. Here, we reveal how such a constraining architecture can still accommodate novel morphs in evolving polymorphisms using the classic polymorphic Batesian mimicry in *Papilio polytes*, whose supergene-like genetic architecture is maintained in a large inversion. We show that rapidly evolving alleles of the conserved gene, *doublesex*, within this inversion underlie the genetic basis of this polymorphism. Using precisely dated phylogeny and breeding experiments, we show that novel adaptive mimetic morphs and underlying alleles evolved in a sequentially dominant manner, undergoing selective sweeps in the mimetic species as predicted under Haldane’s sieve. Furthermore, we discovered that mimetic forms share precise inversion breakpoints, allowing rare exon swaps between the universally dominant and a recessive allele to produce a novel, persistent intermediate phenotype, ultimately facilitating the acquisition of phenotypic novelty. Thus, genetic dominance, selective sweeps, rapid molecular divergence, and rare recombination promote novel forms in this iconic evolving polymorphism, resolving the paradox of phenotypic novelty arising even in highly constrained genetic architectures.

## Introduction

Nearly a hundred years ago, JBS Haldane—one of the architects of Modern Synthesis— predicted that novel adaptive mutations under strong selection will rise rapidly to a high frequency if they are genetically dominant over wild-type^1^. This is because the new, even partially dominant mutations are exposed to selection instantly, rather than suffer potential loss by random genetic drift, even when they initially appear as heterozygotes^1^. This is now known as Haldane’s sieve. In undefended prey species or Batesian mimics, the mutation that confers phenotypic similarity to well-protected aposematic models is under strong selection because it confers high fitness by reducing predation pressure. Thus, this adaptive mutation will spread rapidly under strong selection if it is dominant, as Haldane predicted. Indeed, mimetic forms in polymorphic mimicry in *Papilio* swallowtails are often genetically dominant over non-mimetic forms^2^. Two critical predictions follow from this: (a) mimetic forms that evolve in succession are likely to be sequentially dominant, and (b) the mimicry gene will show signatures of selective sweeps and/or episodic selection because of a few successive mutations that improve mimetic resemblance. However, testing the Haldane’s sieve hypothesis has been difficult in wild populations since the precise evolutionary chronology of mimetic forms in relation to non-mimetic relatives among these species, the dominance relationships among different female forms, and the precise molecular identities of the underlying genes and alleles have been poorly characterised. Likewise, strongly advantageous novel mutations that sweep through populations may completely replace wild-type alleles, making it difficult to estimate their evolutionary origins and the dominance relationships amongst them. On the contrary, selection for balanced polymorphisms in Batesian mimicry may maintain successive beneficial alleles, preserving their evolutionary histories in relation to the original wild-type (*f. cyrus* in case of the *polytes* species group; Fig. 1). This makes mimicry polymorphisms an excellent system to test the predictions of Haldane’s sieve, which we do here by uncovering the missing pieces mentioned above. Finally, we ask how molecular evolution and dominance interact with other genetic aspects to facilitate the evolution of novel forms in an existing polymorphism.

**Figure 1:**
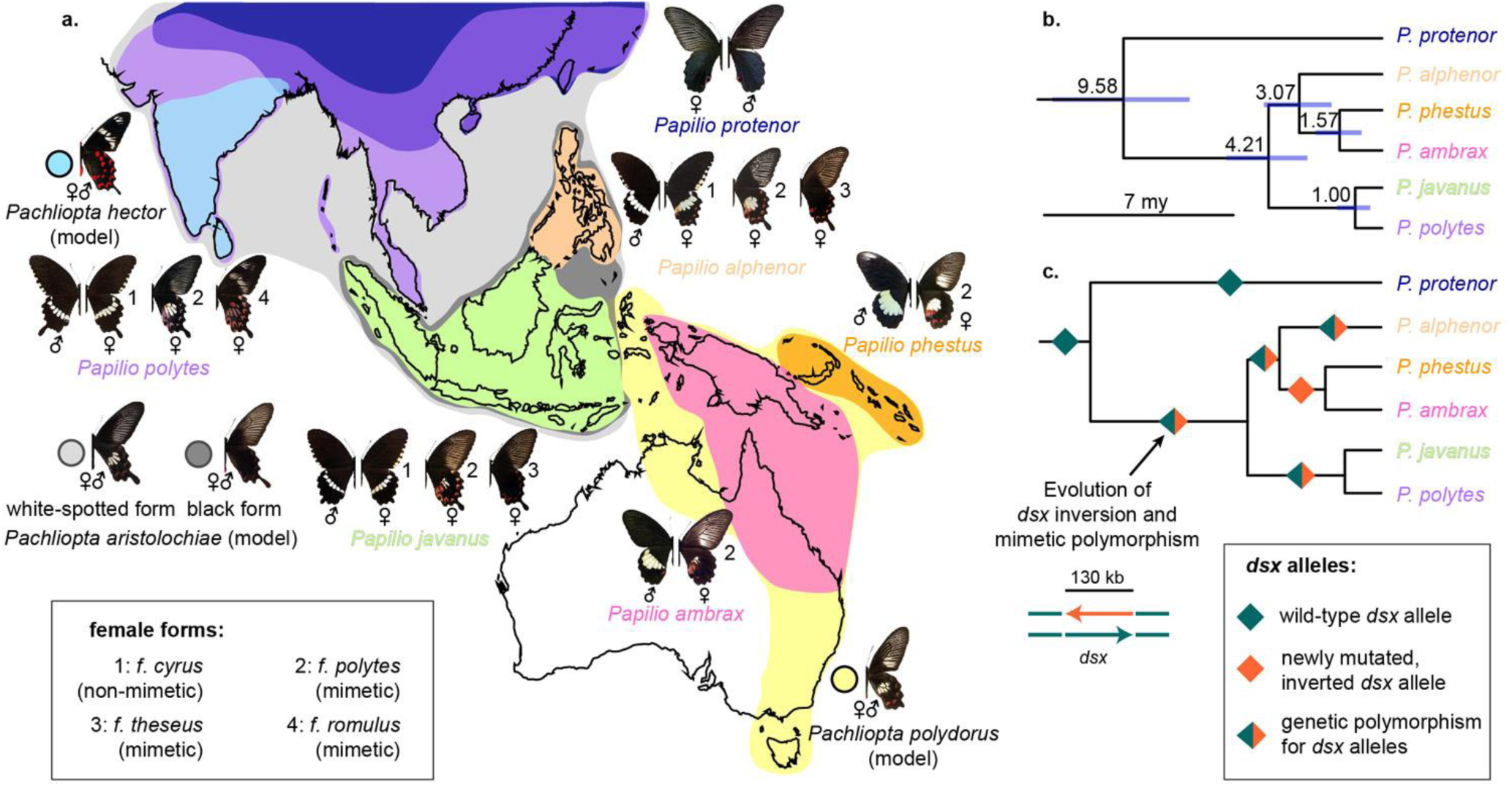
Speciation, wing pattern evolution and mimetic polymorphism in the *Papilio polytes* species group. **a.** Species distributional ranges and wing colour pattern polymorphism in the female forms of the *polytes* species group, along with their Batesian models. Although the minute details of mimetic wing colour patterns and the presence/absence of tails vary across species and populations, the form names are generalised for the purpose of this paper and apply to multiple species. **b.** A secondary fossil-calibrated, dated phylogeny of the *polytes* species group, showing mean with 95% Highest Posterior Density of each split in million years. **c.** Evolution of mimetic polymorphism and *doublesex* inversion in relation to speciation events. Diamonds on branches show fixation or polymorphism of female wing patterns and the evolution of accompanying *dsx* inversion.

*Papilio* swallowtail butterflies exhibiting polymorphic and female-limited mimicry are classic examples of complex polymorphisms^3–5^. As edible Batesian mimics, they derive protection from predators by mimicking toxic, warningly patterned (i.e., aposematic) butterflies. Exemplifying this is the *Papilio polytes* species group, which exhibits four female-limited mimetic morphs or forms (‘*f.*’): *f. cyrus* is male-like and non-mimetic, whereas *f. polytes*, *f. theseus* and *f. romulus* are Batesian mimics of aposematic *Pachliopta* models^3,6^. Female forms *cyrus* and *polytes* occur in several species across the range of the group (Fig. 1a). However, *f. romulus* is endemic to S. Asia and *f. theseus* is endemic to islands of SE Asia, because their aposematic models (*Pachliopta hector* and the black form of *Pachliopta aristolochiae*, respectively) are endemic to those regions^6^ (Fig. 1a). The diversity of these mimetic polymorphisms peak in Sri Lanka and India, where forms *cyrus*, *polytes* and *romulus* co-occur^7^, and in SE Asia where forms *cyrus*, *polytes* and *theseus* co-occur^6^ (Fig. 1a). The female forms in *P. polytes* are autosomally inherited and governed by a single locus, *H* ^6,8^, which was recently identified to be *doublesex* (*dsx*)^9,10^—a developmental master regulator. Although *dsx* controls early developmental sex differentiation in insects, it has been co-opted repeatedly in pupal stages to produce ecologically relevant polymorphisms such as polymorphic mimicry in *P. polytes*, evolving rapidly under positive and pervasive selection^11^. An inversion spanning the entire mimetic *H* allele generates a supergene-like architecture and largely prevents recombination ^9^. How does a single conserved gene such as *dsx* with a constraining genetic architecture promote the remarkable phenotypic diversity observed across lineages in *Papilio*?

## Results and Discussion

### 1. Evolutionary chronology of mimetic forms

We showed recently that *P. polytes* is not a single species, as traditionally believed^6,12^, but a group of three allopatric species in two clades: (a) *P. polytes* (Asian mainland) and its sister *P. javanus* (mainly Sunda Islands), and (ii) *P. alphenor* (the Philippines), sister to the species pair *P. phestus* and *P. ambrax* (Fig. 1a–b)^13^. They show hallmarks of distinct species such as well-supported phylogenetic structure, genome-level divergence, strong assortative mate preference, and postzygotic barriers to hybridization^13^. This discovery reveals the evolutionary history of mimetic polymorphism in this iconic species group, based on the unequal distribution of various female forms across species and populations (Fig. 1). To determine the evolutionary chronology of different female forms along with the underlying mimicry gene inversion among species, we used secondary fossil calibration as previously applied in *Papilio*^14,15^ to date the species phylogeny (Fig. 1b–c, Fig. S1; sample details in Table S1). We estimated that the group is 9.58 million years (my) old (95% highest posterior density (HPD): 12.04 to 7.08 my), with the sexually monomorphic *P. protenor* being basal (Fig. 1b–c, Fig. S2). We used a whole-genome dataset of the *polytes* group as well as inversion breakpoint-specific primers (Table S2) to track the evolutionary history of the *dsx* inversion. The orientation of the *dsx* homologs of outgroups^9,16^ and the basal *P. protenor*^12^ was inferred to be ancestral. Mimicry-related sexual dimorphism, and the *dsx* inversion, is a derived state shared by all descendent species, and is estimated to have evolved between 9.58 my and 4.21 mya (95% HPD: 5.76 to 2.77 my; Fig. 1b–c, Fig. S2). The ancestral orientation of *dsx* and *f. cyrus* were lost, and the *dsx* inversion and mimetic *f. polytes* were fixed, in the ancestor of *P. phestus* and *P. ambrax* (Fig. 1c). However, both mimicry and inversion polymorphisms were retained in *P. alphenor*, *P. javanus* and *P. polytes* (Fig. 1a, c). Females of these three species share the non-mimetic *f. cyrus* and mimetic forms *f. polytes* and *f. theseus*. We included multiple Batesian models of mimetic female forms that occur over their ranges, including all three models from the Indian subcontinent (*Pachliopta aristolochiae, Pachliopta hector* and *Pachliopta pandiyana*) in the time-calibrated phylogenetic reconstruction. The split between *Pachliopta aristolochiae* and *Pachliopta hector* is estimated to have occurred around 0.45 mya (95% HPD: 0.82 to 0.15 my; Fig. S2). The wing pattern of *Pachliopta aristolochiae* is shared among several aposematic *Pachliopta* and related genera that occupy the mainland and islands in the Indo-Australian region, whereas *P. hector* has a unique wing pattern with two white bands on the forewing and no white spots on the hindwing (Fig. 1a). Thus, based both on the occurrence of female *f. romulus* in *P. polytes* alone (split from its sister *P. javanus* at approx. 1 my) and the evolution of the unique wing pattern of the Batesian model *Pachliopta hector* (split from its sister *Pachliopta aristolochiae* at approx. 0.45 my), we infer the evolution of *f. romulus* to be less than 0.45 my (95% HPD: 0.82 to 0.15 my) under a new protective umbrella of the S. Asia-endemic *Pachliopta hector* (Fig. 1b–c, Fig. S2). We tried to find sequences of the *dsx* inversion breakpoints in the genome sequences of all species in the subgenus *Menelaides* but found no sequence matches in any species outside of the *P. polytes* species group. Thus, the mimicry-related inversion found in the *P. polytes* species group appears to be restricted to that group alone. This implies that whatever long-term balancing selection that different *dsx* alleles underlying different female forms of *P. polytes* have experienced also span only the estimated approx. 9.58 my for *f. cyrus* and *f. polytes*, and approx. 1 to 0.45 my or shorter for *f. romulus*. However, smaller inversions in parts of *dsx* appear to be associated with mimicry in species of the *memnon* and *gambrisius* species groups^17^.

### 2. Mimetic polymorphism in *P. polytes* maps to *dsx* alleles and their expression

The genetic basis of forms *cyrus* and *polytes* was earlier mapped to *dsx* alleles^9,10^. It was possible that *f. romulus* of *P. polytes* is regulated by another gene since it involves pigmentation (red spotting and white bands) that is different from *f. polytes* (white spotting). We identified the mimicry locus in *f. romulus* with a genome-wide association study using a *romulus-cyrus* segregating brood. GWAS revealed 20 sites significantly associated with *f. romulus*, all of which mapped to the *dsx* and neighbouring scaffolds that retain synteny with the *dsx*-containing chromosome 25 in *Bombyx mori* (Fig. S3, Table S3). Surprisingly, there were no significant hits within the *dsx* gene, which indicated that reads from *f. romulus* did not map to the *cyrus* allele, perhaps due to an inversion and subsequent genetic divergence. To test this possibility, we sequenced whole genomes of 41 wild-caught *P. polytes* including all three female forms across its geographical range (Fig. S1, Table S1). We found that reads from *romulus dsx* mapped to the mimetic *H* allele of *f. polytes*, and a few reads sparsely mapped to exons of the non-mimetic *h* allele. This indicated that *romulus dsx* occurred in the same inverted orientation as the *H* allele of *f. polytes*, which we now distinguish as *dsx* alleles *H^R^* and *H^P^*, respectively, and *h* referring to the recessive *cyrus* allele. To test whether *H^R^* and *H^P^* have the same *dsx* inversion, we inspected their breakpoints using breakpoint-specific primers (Table S2) in wild-caught as well as lab-bred individuals, including mapping broods. We found that the breakpoints of *H^R^* and *H^P^*were indistinguishable, confirming that the two mimetic alleles have a common evolutionary origin (Fig. S4).

Apart from sharing inversion breakpoints, *H^R^* and *H^P^* alleles were expressed at similar levels in the developing wings of 3-day old female pupae (mimetic wings) compared to those in males (non-mimetic wings) (Fig. S5). This 3-day pupal stage is a critical window for wing colour pattern differentiation of mimetic females in this species group^9,10^. However, isoform expression in the female forms was distinct: of the three known female isoforms of *dsx, dsxF1* was expressed predominantly in *f. romulus* wings compared to *f. polytes* wings (Fig. S5). Moreover, each isoform differs in sequence and structure at the C-terminal end, which potentially alters the DNA-binding ability and facilitates regulation of different downstream effectors by *dsx* to produce the mimetic and non-mimetic wing patterns^10,18^. Thus, evidence from genotype-phenotype mapping (Fig. S3), developmental expression (Fig. S5) and allelic characterisation (see below) shows implies that the *H^R^* allele and its isoforms genetically and developmentally define and developmentally regulate *f. romulus*, even though it shares the same inversion breakpoints, many of the defining SNPs and pupal expression levels of the *H^P^* allele of mimetic *f. polytes*.

Similar to *f. romulus*, the inversion breakpoints of *f. theseus* were identical to that of *f. polytes*, as seen in the alignment of genomic data (Fig. S4). This confirmed that the *dsx* inversion occurred only once and was shared across all the mimetic female forms. However, the *dsx* coding sequence of *f. theseus* was identical to that of *f. polytes* in both *P. javanus* and *P. alphenor* (Fig. 3). This suggests that *f. theseus* may be a developmentally regulated variant of *f. polytes* in which white spots on hindwings are not expressed. This might also explain why *f. polytes* and *f. theseus* show equal dominance^6^. Similar to *f. polytes*, *f. theseus* possibly evolved in the common ancestor of *P. javanus* and *P. alphenor* since it is shared between them (Fig. 1a–c). In a previous study, Zhang *et. al.,* (2017) discovered different non-overlapping sets of *theseus-*specific SNPs in *Papilio polytes* and *Papilio alphenor*. Based on our finding that *Papilio polytes* is a complex of three species, we note that the Zhang *et. al.* study compared *theseus* females from *P. javanus* (sampled from Indonesia) with *polytes* females from *P. polytes* (sampled from Japan). Therefore, what they obtained were species-specific differences and not form-specific differences, which were non-overlapping between the two sets of *theseus* samples. The lack of sequence-level differences between forms *polytes* and *theseus* is encapsulated in our gene tree (Fig. S7), where both forms co-occur on the same branch, whether coding or non-coding *dsx* sequences are considered. Mutations in regulatory elements of *dsx* or other genes involved in wing patterning and pigmentation pathways affecting developmental regulation alone – which do not affect dominance relationships between forms^6^ – may explain the maintenance of *polytes* and *theseus* forms in the island species. This mechanistic basis regarding its origin needs to be confirmed with further developmental manipulations.

### 3. Successive mimetic forms are sequentially dominant

An intriguing difference in the genetic basis of wing patterning in *Papilio* compared to *Heliconius* butterflies is the nature of genetic dominance. *Papilio* usually have a single locus governing polymorphic wing patterns, and morphs are determined by complete dominance in wing pattern inheritance^6,19–21^ whereas in *Heliconius,* several loci govern different aspects of wing patterning, and they often exhibit co-dominance among mimetic forms^21,22^. Complete dominance might have been selected in *Papilio* Batesian mimics to protect mimetic forms from maladaptive intermediates. On the other hand, selection of co-dominance in the aposematic *Heliconius* might have led to novel combinations of wing patterns, which appear to be favoured in this group^23^. To characterise dominance relationships in *P. polytes*, we performed a series of crosses between pure-breeding lines of all three female forms, designed to exploit the known female-limitation of mimicry and achiasmatic oogenesis (Fig. S3d). These crosses showed a clear and complete dominance hierarchy between the three *P. polytes* female form alleles, with *romulus* > *polytes* > *cyrus*, and all segregating broods showing ∼1:1 ratio of dominant and recessive phenotypes, as expected under complete dominance (Table S4–5). These dominance relationships, and the strict allelic inheritance and expression of mimicry in a sex-specific manner, were confirmed by genotyping of parents and female offspring using our form-specific primers (Methods, Table S2). None of the segregating broods showed intermediate or otherwise unusual wing colour phenotypes even when they were grown across different seasons in slightly variable climatic conditions and nutrition regimes, showing that environmental variation or genetic background did not affect the wing patterns (Table S4–5, personal observations). The female forms were thus controlled strictly by their allelic combinations under complete dominance.

The inferred sequence of origins of female forms and its inverse relationship with genetic dominance in *P. polytes* strictly follow the pattern predicted by Haldane: the ancestral, non-mimetic *f. cyrus* that evolved less than 9.58 mya is universally recessive, the next novel mimetic *f. polytes*=*f. theseus* that originated before 4.21 my is dominant over *f. cyrus*, and the most recent *f. romulus* that originated less than 1 mya is universally dominant (Fig. 1). These genetic dominance relationships and sex-limitation of mimicry do not break down in interspecific crosses with monomorphic or polymorphic sister species^6^ – contrary to the pattern observed in *Heliconius numata* where derived dominant alleles also show codominance^24^ *–* indicating that the genetic bases of these two critical adaptations are highly canalised, outweighing the potential impacts of different genomic backgrounds across species.

### 4. *dsx* shows signatures of selective sweeps and episodic selection, with rapidly evolving alleles

Having established that the *dsx* alleles underlie all the mimetic forms including the previously uncharacterised *f. romulus*, and that the successive mimetic forms in the *P. polytes* species group are sequentially dominant, we tested whether *dsx* shows signatures of selective sweeps and/or episodic selection as predicted under Haldane’s sieve. To put this in context, we characterised all the genomic regions that have experienced intense selection in this entire clade by estimating selective sweeps using Raised Accuracy in Sweep Detection (RAiSD). We found that *dsx* is the only gene that has experienced selective sweeps in all five mimetic species in this clade, but not in the basal monomorphic and non-mimetic relative, *P. protenor* (Fig. 2, Table S6). Each species had a different set of *dsx* SNPs showing signatures of selective sweeps (Fig. 2b). All these SNPs were in intronic regions, suggesting that there may be species-specific mutations associated with these SNPs that may have improved mimetic resemblance, which needs to be tested in the future. Further, the Branch-site Unrestricted Statistical Test for Episodic Diversification (BUSTED)^25^ revealed that *dsx* sequences show signatures of episodic diversifying positive selection in the mimetic species (p<0.0001), specifically in codons 142 and 148 (p<0.01) as revealed by Mixed Effects Model of Evolution (MEME)^26^.

**Figure 2:**
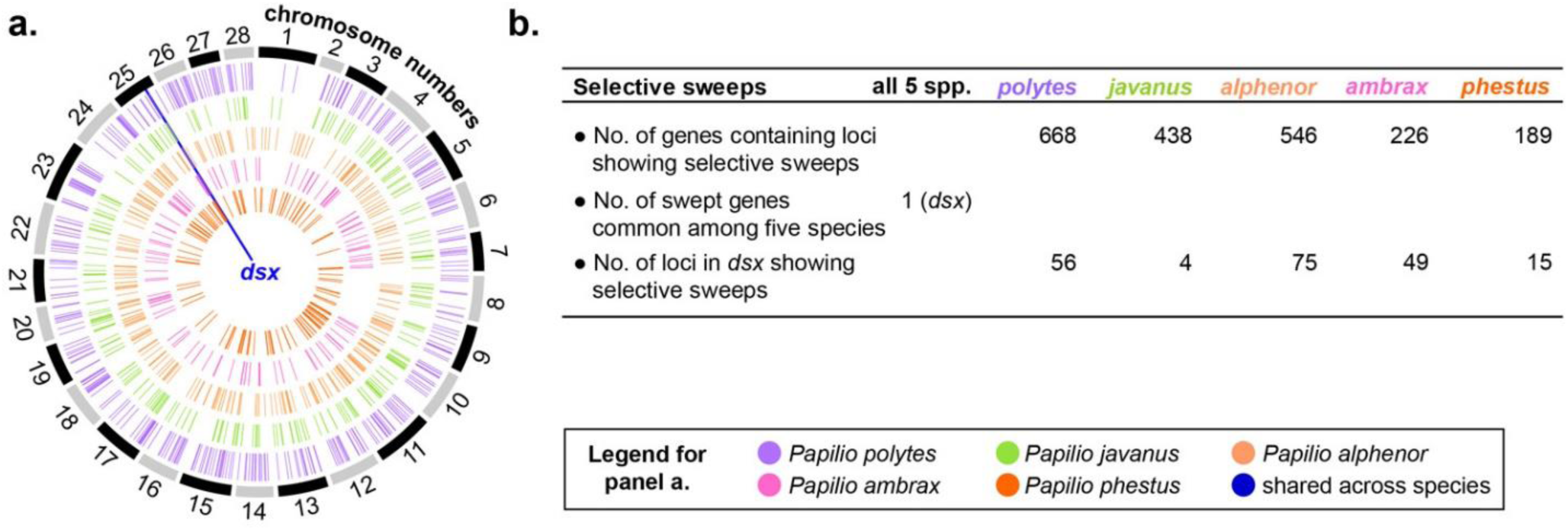
*dsx* is a hotspot of selective sweeps in all the mimetic species. **a.** Annotated genes showing selective sweeps in the five mimetic species. The alternately coloured outer bands indicate chromosomes as defined in the reference *Bombyx mori* genome, whereas colour-coded lines inside represent genes showing signature of selective sweeps in the five species. **b.** Summary of analysis of selective sweeps using Raised Accuracy in Sweep Detection (RAiSD). Note that a single annotated gene may contain multiple loci (SNPs) that show signatures of selective sweeps.

How have the selective sweeps and episodic positive selection influenced molecular diversity and allelic differentiation in *dsx* in the context of mimicry? An analysis of single-nucleotide polymorphisms revealed that the three *dsx* alleles (*H^R^*, *H^P^* and *h*) were highly differentiated: pairwise comparisons of genome sequences of the three female forms revealed high average Fst (0.6 to 1) between variable loci of the three *dsx* alleles (Fig. S6). DNA sequences of *H^R^*, *H^P^* and *h* alleles (*H^R^* n=10; *H^P^* n=35; and *h* n=26) showed a total of 151 fixed substitutions (i.e., 13% of the exonic sequence), which have accumulated in short timeframes with the evolution of the three alleles (Fig. 3–4). Some of these substitutions are in the critically important domain regions of this conserved developmental master regulator, and a few involve amino acid substitutions (Fig. 3). For example, the OD2 domain of *H^R^* has some *romulus-*specific amino acid substitutions which might affect the protein-protein interactions of *dsx* that this domain facilitates (Fig. 3). Interestingly, all these allele-specific substitutions appear to fix early in the evolution of alleles, irrespective of how the female forms are parsed subsequently among species: the *dsx* gene tree showed a well-supported topology in which branches clustered first by colour forms and then by species, rather than the reverse—a pattern also seen in the haplotype network (Fig. S7). The *dsx* gene trees further corroborate the estimated evolutionary sequence of the origin of mimetic forms and the underlying *dsx* alleles (Fig. 1b–c). We did not find any species-specific fixed substitutions or polymorphisms in *dsx* coding sequences of the allelic variants.

**Figure 3:**
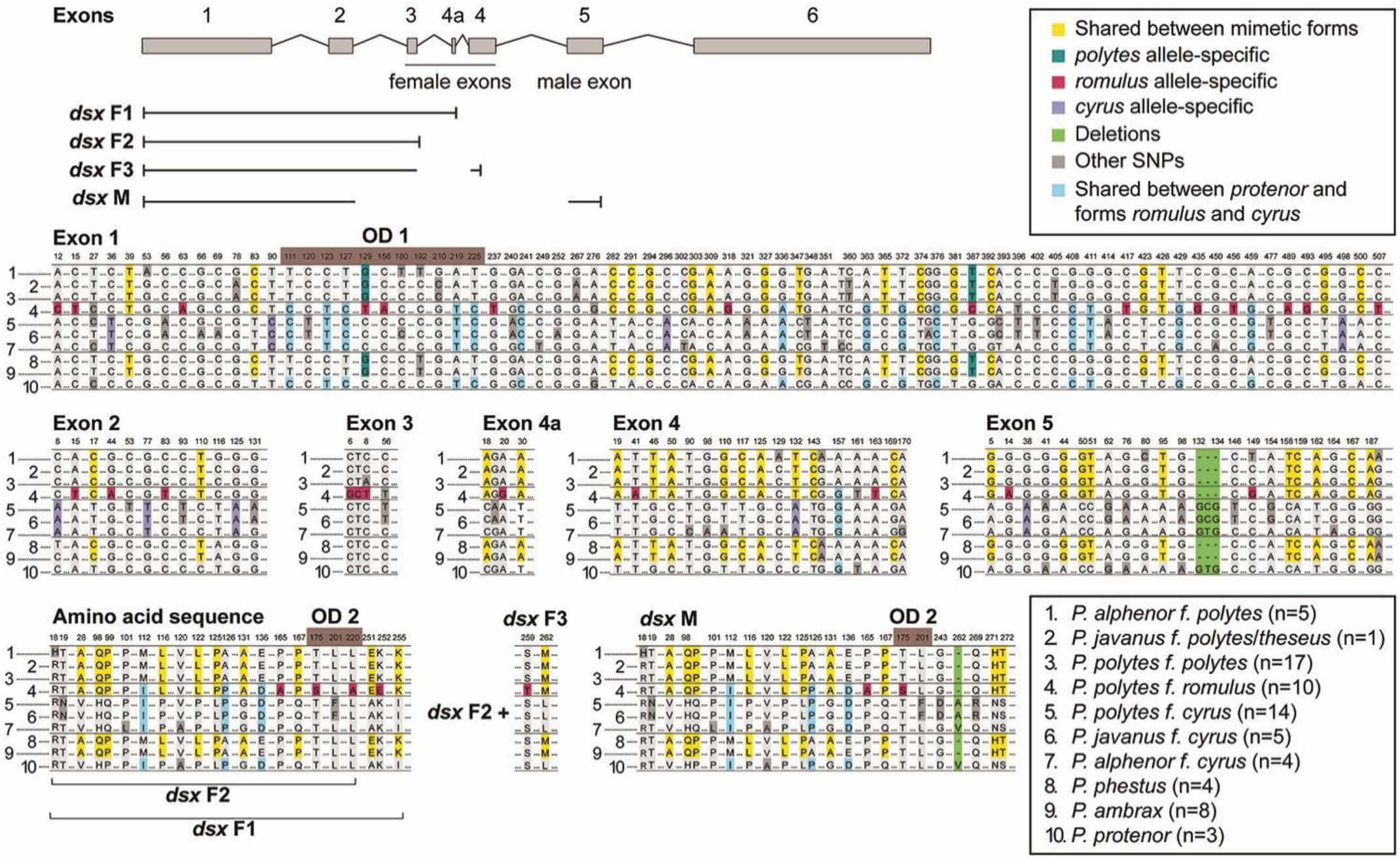
Allelic basis of mimetic polymorphism in the *polytes* species group. Lepidopteran *dsx* comprises of six exons, of which exon 6 is untranslated^27^. It has female- and male-specific transcripts (called *dsx* F and *dsx* M, respectively), with multiple female isoforms (F1, F2 and F3) in the *polytes* group^9,10^. The exon composition of each isoform is depicted at the top. Allele-specific SNPs, their positions on the CDS and protein sequence, and their corresponding amino acids are colour coded. Invariable and non-specific polymorphic sites are represented by dotted lines. The domain regions (OD) are highlighted where feasible. OD2 spans several exonic regions in the DNA sequence where bases cannot easily be numbered because of different lengths of exonic regions across *dsx* isoforms. The amino acid sequence of OD1 is conserved.

**Figure 4:**
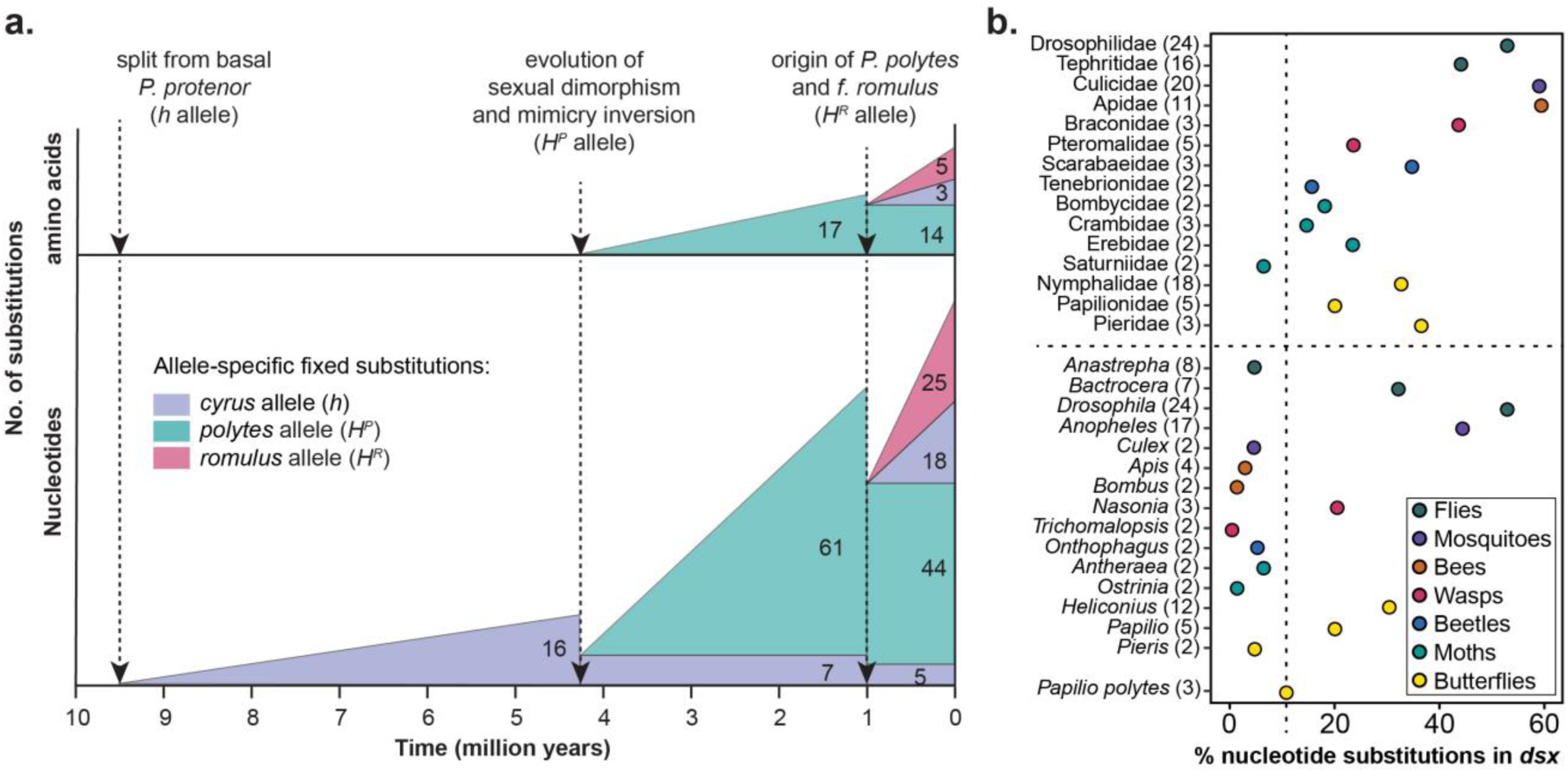
Rapid molecular evolution of mimetic *dsx* alleles in the *polytes* species group. **a.** Allele-specific, fixed substitutions from Fig. 3 are shown for DNA and amino acid sequences (CDS) with respect to the evolutionary timeline of the origin of three *dsx* alleles (a linear increase is assumed for simplicity). The total number of fixed substitutions accumulated in each allele is shown before the new allele evolved. Colour-coded regions that cross evolutionary boundaries (dotted lines) represent the number of fixed substitutions that were inherited in the new allele from the previous allele from which it arose. For example, *h* has no fixed substitutions in the amino acid sequence but 16 in the DNA sequence, of which 7 were inherited in *H^P^*, of which 5 were inherited in *H^R^*. *H^P^* has 61 new, fixed DNA substitutions relative to *h*, of which 44 were inherited by *H^R^*. In addition, *H^R^* and *h* share 18 SNPs that are absent in *H^P^*, and 25 fixed substitutions that are unique to *H^R^*, showing a rapid accumulation of mutations in coding regions of novel mimetic alleles. See Fig. 3 for molecular details and sample sizes. **b.** Percentage of nucleotide substitutions in *dsx* sequence as observed at the genus and family levels (separated by a horizontal dotted line) for four insect orders ^27^. Numbers in parentheses after family/genus names represent the number of species from that group used in this analysis. The three alleles of *dsx* in *P. polytes* alone (vertical dotted line) have more substitutions than the *dsx* sequences within several genera.

The high molecular divergence between the coding regions of the three *dsx* alleles within the *polytes* species group, and especially between the mimetic *H^R^* and *H^P^* alleles of *P. polytes* alone in approx. 1 my, represents an exceptionally high rate of adaptive divergence under positive selection. Indeed, the percentage substitutions observed in *dsx* alleles within this single species is comparable to that across multiple insect genera (Fig. 4b). It is important to note, however, that it is unlikely that every substitution between *H^R^* and *H^P^* alleles is adaptive. It is possible that a considerable number of the fixed differences between *H^R^* and *H^P^* alleles might be a result of hitchhiking associated with strong selective sweeps caused by a few mutations of large fitness effects in the context of mimicry. In any case, this divergence may provide a specific case study and a benchmark for future comparisons of how insect *dsx*, although highly conserved across holometabolous orders, shows rapid differentiation in DNA sequence as well as protein structure when it is under selection for adaptive dimorphisms and polymorphisms in secondary sexual traits ^27^.

### 5. Rare exon swaps produce novel mimetic intermediates

The *dsx* alleles determining different female forms are protected by the inversion or lack of recombination, but how can novel forms evolve in this existing polymorphism? Do they arise strictly from existing forms in a sequential manner via accumulation of substitutions, as suggested above for existing female forms in the *polytes* species group? Or are there other mechanisms, such as evolution in isolation followed by hybridization and introgression^28^, or enhancer shuffling^29^, and differential developmental regulation^30^ by which new forms may be accommodated? A novel female form of *P. polytes* offers a window into this interesting problem. This intermediate form, which we first noticed in nature (Fig. S8, and was also observed by Clarke and Sheppard^6^) and then retrieved in a captive, mixed breeding population (Fig. 5a), had *romulus*-like forewings and *polytes*-like hindwings. We bred this form for over five generations, confirming that it was fertile and genetically stable. We examined the *dsx* sequence of two individuals from this pure-breeding line, which showed that they carried *romulus-*specific SNPs in exon 1, and *polytes-*specific SNPs in exons 2–5 (Fig. 5b–c). This implied that their ancestor had a recombination event in the approx. 35kb intronic region between exon 1 of *H^R^* and exon 2 of *H^P^* alleles. This intermediate phenotype demonstrates that *romulus-*specific SNPs in exon 1 (which contains the DNA-binding domain) may be sufficient to change the phenotype of the forewings but not produce a complete *romulus* phenotype, and that exons 2–5 of *H^P^* could override the dominance of *H^R^*allele to produce *polytes*-like hindwings. The protein-protein interaction domain that lies on other exons may be more critical for patterning hindwings, implying that *dsx* could have form-specific interacting partners, which is also suggested by the *romulus-*specific amino acid substitutions in OD2 domain of *H^R^* (Fig. 3). The 35kb region between exons 1 and 2 may also carry regulatory elements that affect the expression of *dsx*. Although, we do not observe an overall difference in *dsx* expression between these female forms (Fig. S5), the isoform *dsx*F1 seems to exclusively express in *romulus* females. The *dsx* alleles may thus manifest the dominant and recessive phenotypes using a combination of exonic regulation, differential isoform expression, novel downstream targets and interacting partners in each female form^18^.

**Figure 5:**
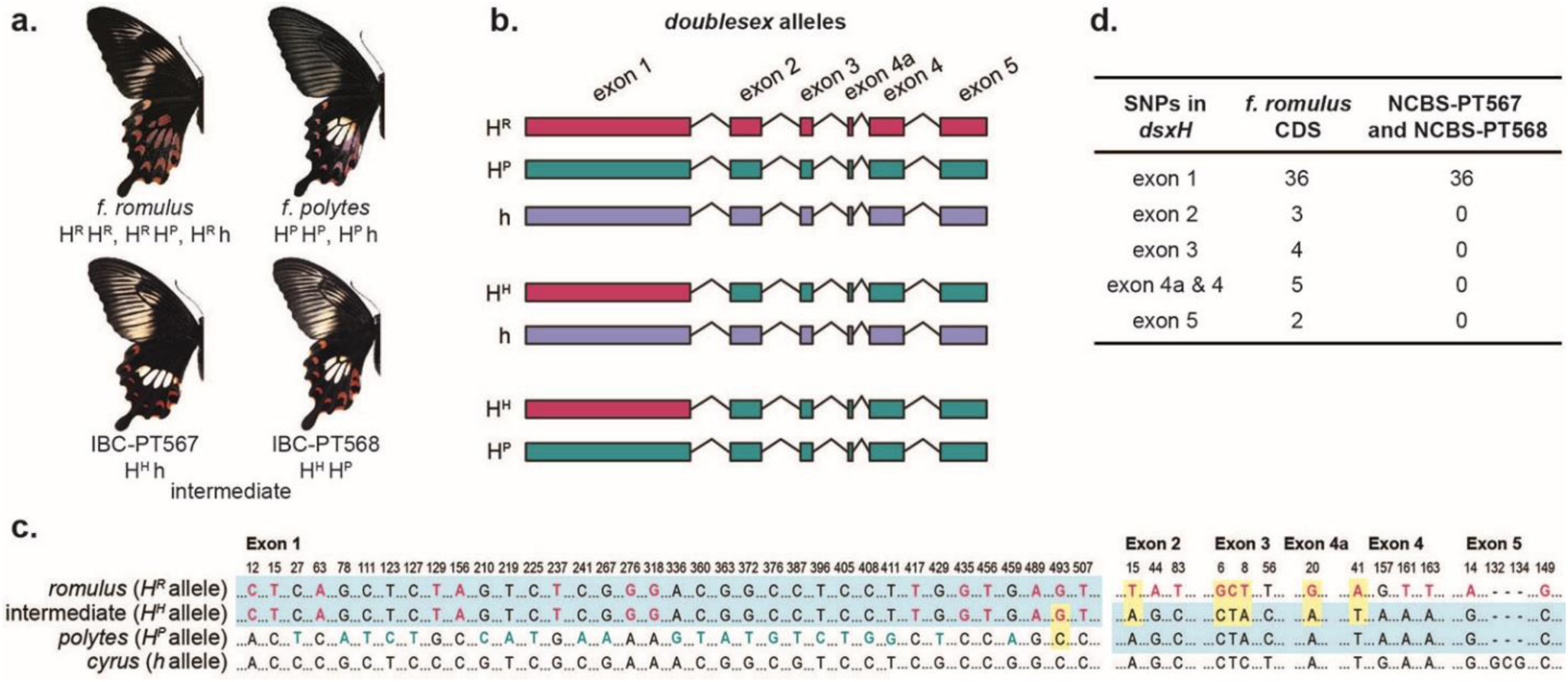
Exon swaps (recombination) between the *H^R^* and *H^P^* alleles produce a novel, rare intermediate phenotype. **a.** Wing patterns and possible *dsx* genotypes of mimetic female forms of *P. polytes*, along with the two females with intermediate wing patterns that we obtained, are shown. Both the intermediates had a crossed over, hybrid *dsx* allele (represented as *H^H^*) with exon 1 of the universally dominant *romulus* allele and exons 2–5 of the second-dominant *polytes* allele. **b.** The three *dsx* alleles have distinctive SNPs in each exon, which are colour coded for the normal phenotypes that they produce. IBC-PT567 was heterozygous for *polytes* and *cyrus* alleles, while IBC-PT568 was homozygous for *polytes* allele, except that both specimens had exon 1 of *H^R^* allele. **c and d.** Exon-specific SNPs of *romulus* and *polytes* alleles in the intermediate IBC-PT567 and IBC-PT568, highlighted in blue. In panel c, *romulus*-specific SNPs are marked in red, and *polytes*-specific SNPs in green. Non-synonymous SNPs are highlighted in yellow.

This persistent intermediate phenotype suggests genetic means, e.g., a rare exon swap, by which novel mimicry phenotypes may be generated. This intermediate phenotype has no existing aposematic model to protect it from predation, so it may be considered at present to be maladaptive. However, such naturally occurring standing variation might become adaptive in other selective landscapes with different distasteful models. Interestingly, other wing pattern combinations, e.g., *polytes*-like forewings and *romulus*-like hindwings, or any other intermediate phenotypes, have never been observed in the field or in the lab. This suggests that the strong genetic architecture and selection together constrain the kind of standing genetic variation in wing patterns observed in this species group.

In light of our overall findings about the evolutionary history of mimetic polymorphism in the *P. polytes* species group and the novel female form just described, it is worth considering the roles of standing genetic variation versus new mutations, potential modification of dominance relationships in relation to Haldane’s sieve, and maintenance of genetic mimetic polymorphisms under long-term balancing selection. There was substantial discussion and speculation about these aspects from the time of the Modern Synthesis to early breeding experiments to probe the inheritance of mimicry^31–33^. With theoretical and molecular genetic advances, a much more nuanced hypothesis has developed. The consensus seemed to be that for Batesian mimicry to evolve, an initial mutation would be required that establishes partial mimetic resemblance to an aposematic model. Due to the complexity associated with wing pattern phenotypes, it is likely that additional subsequent mutations would be necessary to improve mimetic resemblance, and these were termed “modifiers”^34,35^. A modifier could add to mimetic resemblance itself, make it partially or completely dominant, and/or reduce recombination to ensure that the mutations associated with mimicry did not break down and were inherited together. Therefore, modifiers could encompass an array of mutations ranging from point mutations that alter gene function to large inversions and rearrangements that could give rise to supergene architecture. The mimicry “gene” could subsequently either go to fixation, or the mimetic and non-mimetic allelic combinations could be maintained by balancing selection in polymorphic species under negative frequency dependence. In *Papilio polytes*, it was assumed that mimetic polymorphism is maintained by long-term balancing selection on the underlying allelic variation at the mimicry gene *dsx*, as HKA tests showed high genetic diversity at this locus^10,34^. Our work defines the timescales of these long-term balancing selection events to be up to 9.58 my and 1 my for the mimetic forms in the *P. polytes* species group (Fig. 1 and 4, Section 1 of Results and Discussion). Additionally, our analysis revealed that *dsx* has experienced selective sweeps in the mimetic species of the *P. polytes* species group but not in the basal non-mimetic *P. protenor*, implying that mimetic alleles experienced a rapid increase in frequency in the population. This contrast of high genetic diversity and signatures of sweep could co-occur if the mimetic allele shared among the species swept through the population under a soft sweep between 9.58 my and 4.21 mya, before the mimetic species split from each other. The selective sweeps likely caused moderate levels of genetic hitchhiking, and the alleles subsequently accumulated mutations due to lack of recombination and rapid molecular evolution at this locus^11,12^, resulting in high genetic diversity at the locus with some remnants of a sweep.

The origin of the first mutation causing mimetic resemblance may affect the spread of this trait through the population. While this may have come from existing standing genetic variation or new mutations, the latter has largely been implicated in mimicry ^31–33^. The *dsx* gene lies within a specific inversion that seems to be unique to the *P. polytes* species group as we have found no matching inversion breakpoint sequences outside of the *polytes* species group. Yet, *dsx* seems to have been repeatedly recruited in mimetic resemblance in other closely related groups as well, however, with other unique allelic combinations and independent smaller inversions^16,17^. Therefore, along with our genetic analysis of allelic variations and selective sweeps, current evidence appears to support Fisher, Sheppard and Ford’s speculations that mimicry in the *P. polytes* species group may have evolved in large part from novel mutations at *dsx*, many of which apparently sweeping through the populations, rather than from standing genetic variation that was shared between *polytes* and *memnon* species groups. Under this scenario, the predictions of Haldane’s sieve would be valid and directly determined by the dominance of new mutations^36^.

The more challenging aspect of Haldane’s sieve is to identify whether a novel mutation contributing to mimetic resemblance is partially or completely dominant at the point of origin, or it may subsequently become dominant under selection. Under Batesian and Müllerian mimicry, evolution of dominance modifiers may allow an initial recessive mutation to become partially or completely dominant, thereby facilitating its spread through the population^37–39^. It is currently unclear what may constitute a dominance modifier in the context of mimetic resemblance, for example, a regulatory element may alter the expression levels of an allele or an epistatic interaction may alter the dominance of a mutation^40^. However, in terms of evolutionary dynamics under Haldane’s sieve, the spread and the subsequent evolution of this mimicry mutation would be impacted by the point of dominance, not the origin. It is currently methodologically challenging to deduce whether dominance arose with the first mutation leading to a partial mimetic resemblance or evolved subsequently as mimicry was perfected with additional mutations^31,32^, and how much fitness benefit successive mutations offered^41^. In the future, a combination of genetic and developmental manipulations may perhaps be able to throw light on these aspects.

Our recent finding that *P. polytes* is a complex of three species^13^ makes it necessary to reevaluate previous genetic studies that assumed *javanus* and *alphenor* to be subspecies of *P. polytes*. Clarke and Sheppard’s characterisation of inheritance of the female forms was based on interspecific hybrids between the three species rather than segregating broods of the same species^6^. The first genome-wide mapping and developmental expression studies of mimicry polymorphism were done in the Philippine *P. alphenor*, not in *P. polytes*^10^. Subsequent reference genome and developmental studies found inversion breakpoints and differential expression of *dsx* largely in the Japanese populations of *P. polytes*^9,42^. Previous studies determined the evolutionary history of mimetic forms in this group based on samples from different species, which has possibly led to incorrect conclusions regarding the origin of *f. theseus*^12^. However, conclusions regarding the developmental genetic basis of mimicry from the present and previous genetic studies on *P. polytes* and *P. alphenor* are not contradictory, showing that: (a) the developmental genetic basis of mimicry and female polymorphism is shared between both the species, and (b) *dsx* controls mimetic polymorphism by means of: (i) an inversion—which predates the diversification of all mimetic species in the group—that isolates mimetic from non-mimetic alleles, (ii) allelic variation and differential gene expression across males/*f. cyrus* and mimetic females, producing sex-limitation of the mimetic polymorphism, and (iii) tissue-specific isoform expression across developmental stages to accommodate novel adult polymorphisms that are regulated in late development^9,10,42^.

Our work contains extensive sampling of the *polytes* species group, including dozens of specimens of all known female forms from mainland Asia, specifically South Asia where mimetic polymorphism is most striking, and several island groups that were not sampled in previous studies. This extensive sampling and our reconstruction of the evolutionary history of mimetic polymorphism in this group, the origin of each female form, the dominance relationships between alleles of *dsx*, and molecular evolution *dsx* in relation to mimetic polymorphism, provided unparalleled insights into the evolutionary and genomic history of this entire species group that substantially contrast with previous studies^6,12,13^. We have characterised allele-specific mutations across the *dsx* gene, which could be used for functional assessment and identification of key mutations that affect mimetic phenotypes. We demonstrate that the female forms in the *polytes* species group have evolved in a step-wise fashion through Haldane’s sieve— each successive novel mimetic form having complete dominance over pre-existing forms. Further, we find that novel wing phenotypes could arise from rearrangements in existing form-specific alleles, through exon swaps, and that different exons of the same gene could regulate colour patterns on the forewing and hindwing. Our work also sheds light on how dominance may interact with other genetic and population genetic phenomena, such as recombination and selective sweeps, to influence evolutionary trajectories of diversification of lifeforms.

## Materials and Methods

### Specimen collection, genome re-sequencing and SNP calling

We preserved wild-caught *P. polytes* and *P. protenor* in 100% ethanol across the species range (Fig. S1, Table S1). We also obtained *P. alphenor* and *P. javanus* from native commercial butterfly breeding facilities in the Philippines and Java, respectively, and preserved them in 100% ethanol. Sample details are in Table S1. We extracted DNA from thoracic muscle of preserved samples using QIAGEN DNeasy blood and tissue kit, quantified the extracted DNA using Qubit fluorometric quantification, and prepared libraries using Illumina TruSeq DNA PCR-free library preparation kit. We used a 2x100 PE run on Illumina HiSeq 2500 for re-sequencing genomes of our *Papilio* samples. We also downloaded available genome sequences from the SRA database for *P. alphenor, P. javanus, P. phestus, P. ambrax* and *P. protenor*. We checked the quality of raw reads, aligned them to the *Papilio polytes* reference genome (Ppol_1.0, genome version 1.0, annotation version 1.0) from NCBI^9^, using BWA aligner^43^, and called SNPs using the GATK pipeline^44^. We processed the alignment files to mark and remove duplicates, merge files from the same sample, performed indel realignment, called haplotypes, and combined the resulting VCF files for genotype calling. We filtered the resulting SNPs using the recommended parameters in the GATK manual. The resulting dataset contained samples that had an average coverage per sample of 9X across the genome; and 13X and 8X across the mimetic and non-mimetic *doublesex* scaffolds respectively. Exons of the *dsx* gene showed nearly complete coverage and higher depth compared to introns. Post filtering of called SNPs, we obtained a set of 27,965,662 SNPs for the *polytes* species group. We extracted *dsx* CDS from individual alignments using bcftools-1.6 mpileup^45^. We aligned exons manually and annotated allele-specific SNPs to generate the SNP map for *dsx* in Extended Fig. 3. We identified sites that were fixed in all samples of each female form to generate consensus CDS for each *dsx* allele. We also sequenced genomes of two intermediate female individuals from a pure-breeding line, aligned and processed them as given above, and manually annotated SNPs to find out their exonic composition (Fig. 5). We performed nucleotide and protein alignments in MEGA X^46^ and visualised them in Jalview^47^.

### Reconstruction of species phylogeny, gene trees, and haplotype network

We extracted CDS and non-coding sequence of *dsx* from the genome re-sequencing data using bcftools-1.6 mpileup^45^. We aligned sequences with Muscle aligner^48^ in MEGA X, using codon alignment for the *dsx* coding sequence. We used PartitionFinder 2 to find appropriate models of evolution for each set of sequences: F81+G, GTR+I and JC for the *dsx* CDS dataset, GTR+I+G for the *dsx* non-coding dataset. We reconstructed *dsx* gene trees using MrBayes 3.2.7a with the same parameters as mentioned above. We viewed all phylogenies using FigTree v1.4.3^49^. We filled gaps in the CDS extracted from individual alignments using exon-specific PCRs, followed by Sanger sequencing. Primer details are provided in Table S2. We used popart-1.7^50^ to generate a haplotype network of *dsx* alleles using TCS method (Fig. S7).

### Estimation of divergence times

We acquired sequence data for four mitochondrial markers (cytochrome c oxidase subunit-I, tRNA leucine, cytochrome c oxidase subunit-II and 16S) and two nuclear markers (elongation factor-I alpha and wingless) from a broad range of Papilionidae species (Condamine et al 2023^15^ and Joshi and Kunte Biorxiv^51^) to estimate the time of origin of the mimics and models. We aligned the sequences using MAFFT v7.475^52^ and visually verified the accuracy of the alignments using MEGA X^46^. We used PartitionFinder 2.1.1^53^ to identify the best-fitting partitioning scheme and corresponding models of evolution. We used BEAST v2.7.4^54^ for generating the time-calibrated phylogeny. We applied an optimized relaxed clock model and a birth-death model as molecular clock and tree priors, respectively. For time calibration, we used three primary calibration points (the crown ages of (1) Papilionidae, between 47.80 and 150 Ma, (2) Parnassiinae, between 23.03 and 150 Ma, and (3) Luehdorfiini, between 5.33 and 150 Ma) and 10 secondary calibration points (the crown ages of (1) Papilioninae, between 34.40 and 62.90 Ma, (2) Leptocircini, between 26.60 and 49.90 Ma, (3) Troidini, between 26.90 and 50.40 Ma, (4) Papilionini, between 27.50 and 50.90 Ma, (5) Old World *Papilio*, between 21.52 and 33.5 Ma, (6) subgenus *Heraclides*, between 16.07 and 27.48 Ma, (7) subgenus *Araminta*, between 9.89 and 18.61 Ma, (8) subgenus *Papilio*, between 7.01 and 13.85 Ma, (9) subgenera *Achillides*, *Princeps*, and *Menelaides*, between 15.57 and 24.39 Ma, and (10) subgenus *Menelaides*, between 10.90 and 17.65 Ma) with uniform priors^15^. We ran six independent MCMC chains for 150 million generations and sampled every 15,000 generation. We performed the phylogenetic analyses on the CIPRES web server (http://www.phylo.org/index.php/portal). We combined the results using LogCombiner after removing 25% of the samples as burnin and ensured convergence by evaluating effective sample sizes of relevant parameters on Tracer v1.7.1^55^. We constructed the maximum clade credibility (MCC) tree using TreeAnnotator after removing 25% of the samples as burnin. Divergence times estimated by our work were similar to those estimated in previous studies, although we included many more *Menelaides* samples and additional outgroups to estimate node ages of models and mimics in these mimicry relationships.

### Estimation of signatures of selection in *P. polytes, P. javanus* and *P. alphenor*

To identify loci showing signatures of selection in the genomic dataset, we filtered the SNP dataset (total 27 million SNPs) by species and calculated mu statistic using Raised Accuracy in Sweep Detection (RAiSD) v.2.5^56^. We used a conservative cutoff (>99.5%) of the µ statistic to identify loci experiencing selective sweeps. To minimise the possibility of false positives, we randomly resampled SNPs from a subset of samples to calculate the false positive rate threshold in RAiSD. The resulting threshold was lower than the 99.5% cutoff we had used, so it did not eliminate any resulting hits. The number of identified SNPs experiencing selective sweeps that were included within annotated genetic elements and those that were in the intergenic regions lacking annotation features are given elsewhere^13^. The identities of annotated loci experiencing selective sweeps among the species are listed in Table S6. We mapped the genes experiencing selective sweeps in each species and common across the five mimetic species to the *Bombyx mori* genome^57^ using BLAST+ to identify their chromosomal locations. We used Circos-0.69-9^58^ to map these genes across chromosomes for each species, assuming synteny between *P. polytes* and *Bombyx mori* genomes.

We used MEME^26^ to estimate the sites subjected to episodic positive or diversifying selection (Fig. 2) between *dsx* alleles. We used BUSTED^25^ to test for positive selection by asking whether *dsx* has experienced positive selection in at least one site in the allelic variants. We implemented MEME and BUSTED in HyPhy 2.5^59^ using our *dsx* gene tree of coding sequences (Fig. S7).

### Cross design and dominance hierarchy

We used mated wild-caught females from NCBS campus to start *P. polytes* greenhouse populations, which we maintained in large (2x2x2m) cages. We raised larval broods at approx. 28±4°C, on a diet of lemon (*Citrus sp.*) and curry plants (*Murraya koenigii*) and maintained the adults on Birds Choice™ butterfly nectar. We generated mapping broods (design described in Fig. S3d) by separating mating pairs from the population and letting the mated females lay eggs in individual small (0.6x0.6x1m) cages. Because of achiasmatic oogenesis, we then back-crossed the heterozygous male offspring normally with homozygous recessive females from the greenhouse populations. For GWAS, we generated a total of six segregating broods (1:1 mimetic:non-mimetic female progeny), three each segregating for *polytes/cyrus* and *romulus/cyrus* phenotypes. Among these, we sequenced the parents and 50 female offspring of each form from a *romulus-cyrus* segregating brood that produced 426 offspring. We used these six broods as well as similarly generated *romulus-polytes* segregating broods for genotyping with form-specific primers (Table S2) to establish the dominance hierarchy (TableS4–5). Among 290 individuals genotyped from 10 mapping broods, we observed no cross-reactivity of primers between forms, indicating that these SNPs are indeed form-specific, and that the colour forms are inherited and expressed in a sex-specific manner.

### Genome-wide association study

We used a reduced representation genotyping-by-sequencing (GBS) approach to sequence offspring of the largest *romulus-cyrus* segregating brood, with a standardised protocol for generating a marker set using *Pst*I and *Mlu*CI*-*mediated shearing of DNA, followed by library preparation and sequencing on an Illumina platform using 75 bp SE module (Genotypic Technologies Pvt. Ltd.). We used a TASSEL GBS pipeline with the *Papilio polytes* reference genome for SNP calling^60^ and tested the association of SNPs with female forms using a dominant model. We found very few reads from *romulus* individuals mapping to the non-mimetic *cyrus* allele of *dsx* from the reference genome. This may be due to low representation and coverage of the *dsx* gene with the GBS method, or due to mismatch between the highly divergent *romulus* and *cyrus dsx* sequences. We used Bonferroni correction to account for multiple comparisons to call significant associations. We used BLAST on NCBI and KaikoBase platforms to annotate the sites that associated significantly with *f. romulus* (Fig. S3).

### Testing genetic divergence between the *dsx* alleles

To find genetic divergence between *dsx* sequences from different samples, we grouped our dataset by female forms, and extracted SNP data for the *dsx* mimetic and non-mimetic scaffolds. We calculated Weir-Cockerham’s Fst for the three female forms using vcftools 0.1.13^61^ (Fig. S6). To compare substitution rates between *dsx* alleles in *P. polytes* and other Lepidoptera, we used *dsx* alignments from Baral et. al., 2019^27^, and identified substitutions in *dsx* sequences at the level of genera and families (Fig. 4b).

### Breakpoint characterization for *f. romulus*

To identify potential breakpoints of *H^R^*, we designed primers flanking breakpoints of *H^P^* ^9^. We performed PCRs with lab-bred and wild-caught samples of all three female forms, cleaned the PCR products with ExoSAP-IT™ cleanup reagent and sequenced them using Sanger sequencing. We then manually aligned the sequences of the left and right breakpoints in MEGA X and annotated the alignment using Jalview (Fig. S4).

### qPCR for *dsx* expression

We extracted RNA from 3-day pupal tissue using phenol-chloroform extraction method. We synthesized cDNA using ProtoScript II First Strand cDNA Synthesis Kit (New England Biolabs) and performed qPCR with existing primers for *dsx* and its isoforms^9^ using *RPL3* as an endogenous control (our *RPL3* primers given in Table S2, Fig. S5).

## Acknowledgements

We thank Mark Kirkpatrick and Magnus Nordborg for insightful discussions on population genetics, recombination and the evolution of protected polymorphisms; Anuj Jain and Adam M. Cotton for contributing samples; NCBS Sequencing Facility, AgriGenome, and Genotypic for sequencing; Bhavya Dharmaraaj and Tulsamma for maintenance of greenhouse populations and broods of *Papilio polytes*; NCBS Greenhouse Facility for breeding butterflies, and Biodiversity Lab Research Collections for sample preservation. Wild-caught samples were obtained largely from the NCBS campus/field stations and private lands, and from wildlife sanctuaries and national parks in India under the following research and collection permits issued by the state forest departments in Karnataka (permit no. 227/2014-2015 dated 2015/04/16), West Bengal (permit no. 2115(9)/WL/4K-1/13/BL41, dated 2013/11/06; and permit no. 1107/42/2W-705/18, dated 2018/05/07), Sikkim (dated 2011/03/21), and Meghalaya (permit no. FWC/G/173/Pt-II/474-83, dated 2014/05/27), for which we thank the Principal Chief Conservator of Forest, Deputy Conservators of Forest, Wildlife Wardens and field officers of those states.

## Author contributions

RD and KK designed research; RD and SB generated and analysed genome and genotype sequence datasets; RD and MK performed selection analyses; AG and SB performed phylogenetic analyses; AG performed species delimitation analyses; KK conceived and directed the project; KK and RD wrote the paper.

## Data Availability Statement

All the sequence data used in Fig. 1, 3, Table S1, Fig. S3–7 will be submitted to NCBI, and phylogenetic trees (Fig. 1, Fig. S2, S7) will be submitted to Dryad, after the manuscript is initially accepted. Accession numbers will be provided in the final manuscript after initial acceptance.

## Additional Information

***Supplementary Information*** is available for this paper.

***Correspondence and requests for materials*** should be addressed to

## Supplementary Information

**Table S1:** Sample details of wild-caught butterflies used in the study. See Excel file, TableS1_SampleDetails_PapilioGroup_2019-04-30.xlsx.

**Table S2:**
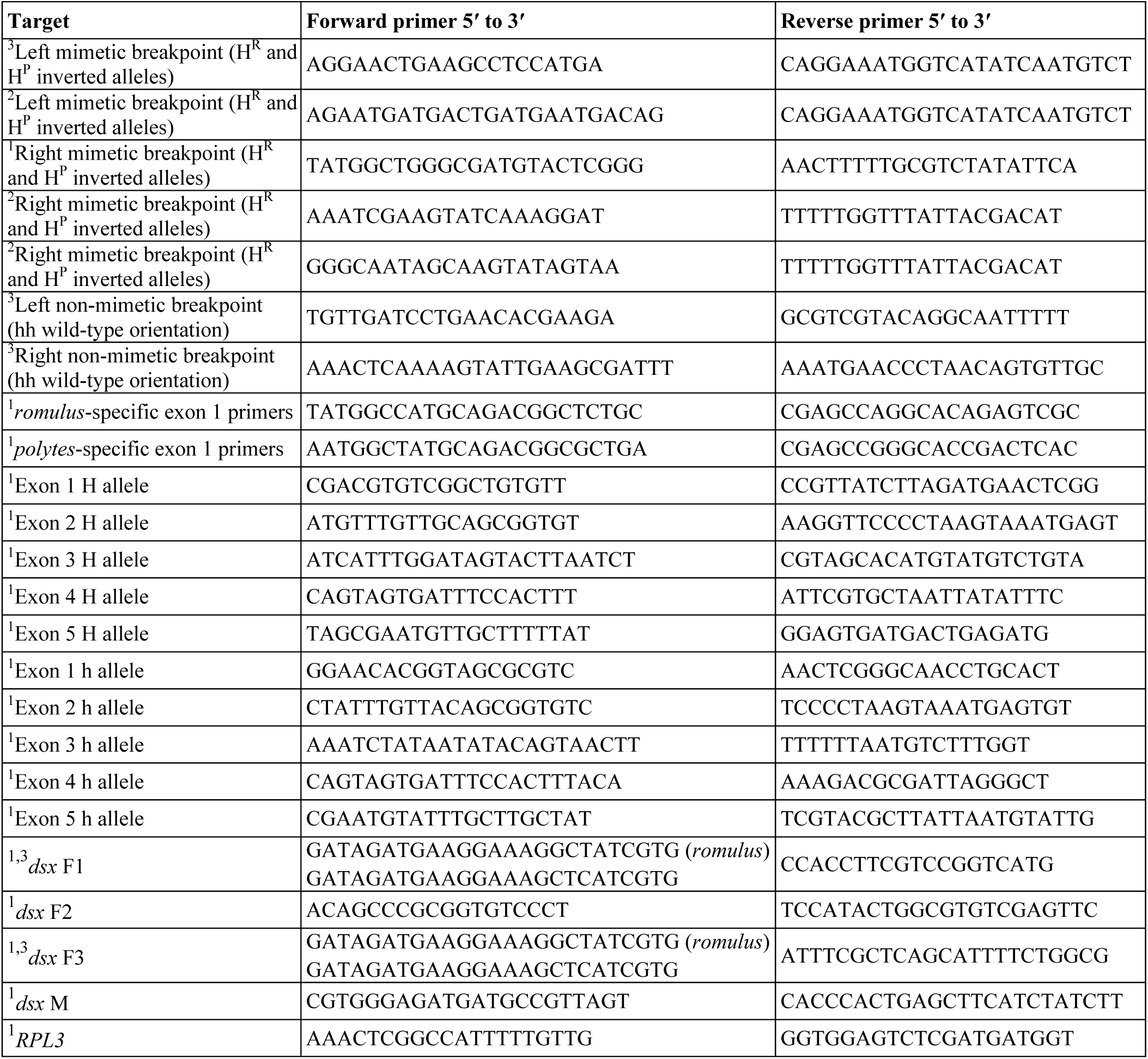
Primers used in this study. Breakpoint- and exon-specific primers distinguish between the three female forms (considering *polytes* and *theseus* forms to be genetically indistinguishable) in the *polytes* species group. ^1^=primers designed for this study, ^2^=primers previously published^8^ but that failed in our PCRs, and ^3^=primers previously published^8^ that also worked in our samples.

**Table S3:** Details of the significant sites associated with the mimetic *f. romulus* in the GWAS.

| SNP | -log<sub>10</sub> (p value) | Locus | Gene | Mapped to *Bombyx mori* chromosome no. |
| --- | --- | --- | --- | --- |
| Sgi.908437033.ref.NW_013528582.1._2192979 | 1.25E-24 | LOC106110479 | putative epidermal cell surface receptor | 25 |
| Sgi.908437033.ref.NW_013528582.1._1181012 | 4.25E-22 | LOC106110489 | phosphofurin acidic cluster sorting protein 2 like | 25 |
| Sgi.908437033.ref.NW_013528582.1._1352228 | 2.75E-19 | LOC106110471 | papilin | 25 |
| Sgi.908463965.ref.NW_013525515.1._13588 | 3.70E-18 | LOC106104332 | ubiquitin carboxyl-terminal hydrolase 34 | 25 |
| Sgi.908466945.ref.NW_013525180.1._33703 | 4.63E-18 | LOC106100995 | cilia and flagella-associated protein 36 | 25 |
| Sgi.908464090.ref.NW_013525504.1._388409 | 1.15E-17 | LOC106103840 | sodium/potassium transporting ATPase subunit | 25 |
| Sgi.908466945.ref.NW_013525180.1._33738 | 7.29E-17 | LOC106100995 | cilia and flagella-associated protein 36 | 25 |
| Sgi.908437033.ref.NW_013528582.1._394916 | 9.47E-16 | LOC106110409 | nephrin-like | 25 |
| Sgi.908464090.ref.NW_013525504.1._440516 | 9.88E-14 | LOC106103855 | Dihydroxyacetone phosphate acyltransferase | 25 |
| Sgi.908466945.ref.NW_013525180.1._932743 | 9.55E-13 | LOC106100916 | ribosome binding protein 1 | 25 |
| Sgi.908466945.ref.NW_013525180.1._419975 | 9.60E-13 | LOC106101076 | uncharacterised locus | 25 |
| Sgi.908466945.ref.NW_013525180.1._532434 | 6.48E-12 | LOC106101139 | probable ATP-dependent RNA helicase spindle E | 25 |
| Sgi.908437033.ref.NW_013528582.1._1346921 | 5.46E-10 | LOC106110471 | papilin | 25 |
| Sgi.908437033.ref.NW_013528582.1._1180988 | 1.74E-08 | LOC106110489 | phosphofurin acidic cluster sorting protein 2 like | 25 |
| Sgi.908437033.ref.NW_013528582.1._1325088 | 2.10E-08 | LOC106110419 | baculoviral IAP repeat containing protein 6 | 25 |
| Sgi.908466945.ref.NW_013525180.1._1924162 | 2.82E-08 | LOC106101077 | serine/threonine-protein kinase D1 | 25 |
| Sgi.908437033.ref.NW_013528582.1._2192947 | 4.82E-08 | LOC106110479 | putative epidermal cell surface receptor | 25 |
| Sgi.908437033.ref.NW_013528582.1._2192954 | 4.82E-08 | LOC106110479 | putative epidermal cell surface receptor | 25 |
| Sgi.908437033.ref.NW_013528582.1._1325082 | 7.03E-08 | LOC106110419 | baculoviral IAP repeat containing protein 6 | 25 |
| Sgi.908466945.ref.NW_013525180.1._1452145 | 1.70E-07 | LOC106101073 | autophagy related protein 2 homolog A | 25 |

**Table S4:**
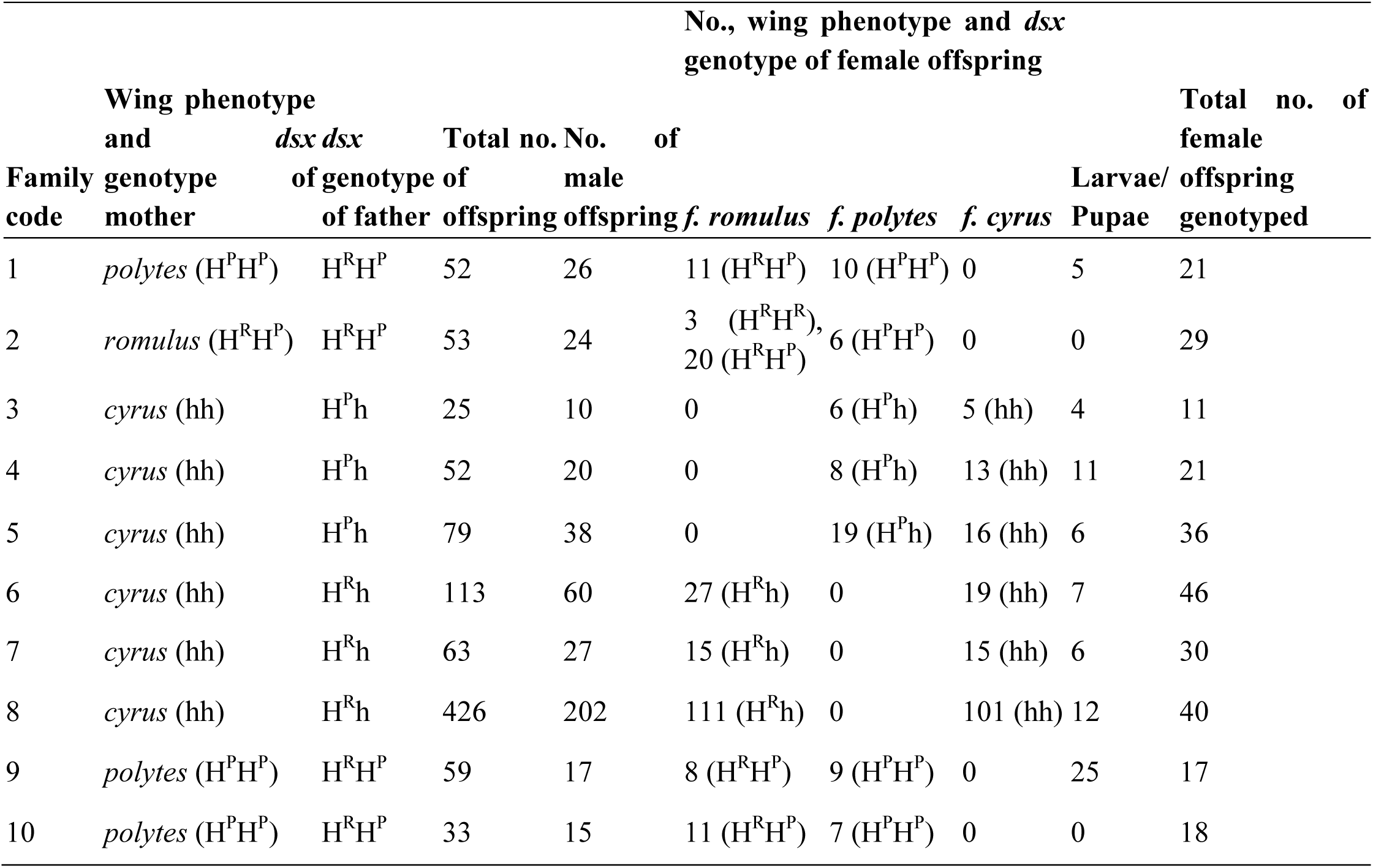
Dominance hierarchy of female forms in *P. polytes*. Details of crosses with various phenotypic and genotypic combinations of parents, and the offspring produced, are shown. The three *dsx* alleles that produce three female forms are listed as *H^R^* (mimetic *romulus* allele that is universally dominant), *H^P^* (mimetic *polytes* allele that is recessive to *romulus* but dominant over *cyrus*), and *h* (non-mimetic *cyrus* allele that is universally recessive). The column “Larvae/Pupae” shows the number of individuals that died in early stages, so their sex and form were unknown. Only female offspring were genotyped since the polymorphic phenotype manifests only in females where genotypes and phenotypes can be correlated. Genotypes were identified using allele-specific primers (Table S2). Additional details of the mapping broods and genotyping (including PCR gel images) are provided in Table S3, and phenotypes are depicted in Fig. 1.

**Table S5:** Mapping brood details, phenotypes and gel images used for testing the dominance hierarchy of the three female forms in *P. polytes*. See Excel file, TableS3_DominanceHeirarchy_GenotypingMappingBroods_2018-09-12.xlsx.

**Table S6:** Loci experiencing selective sweep. Annotated results of RAiSD analysis for selective sweeps. See Excel file, TableS4_LociUnderSelection_RAiSD_GeneAnnotations.xlsx.

**Figure S1:**
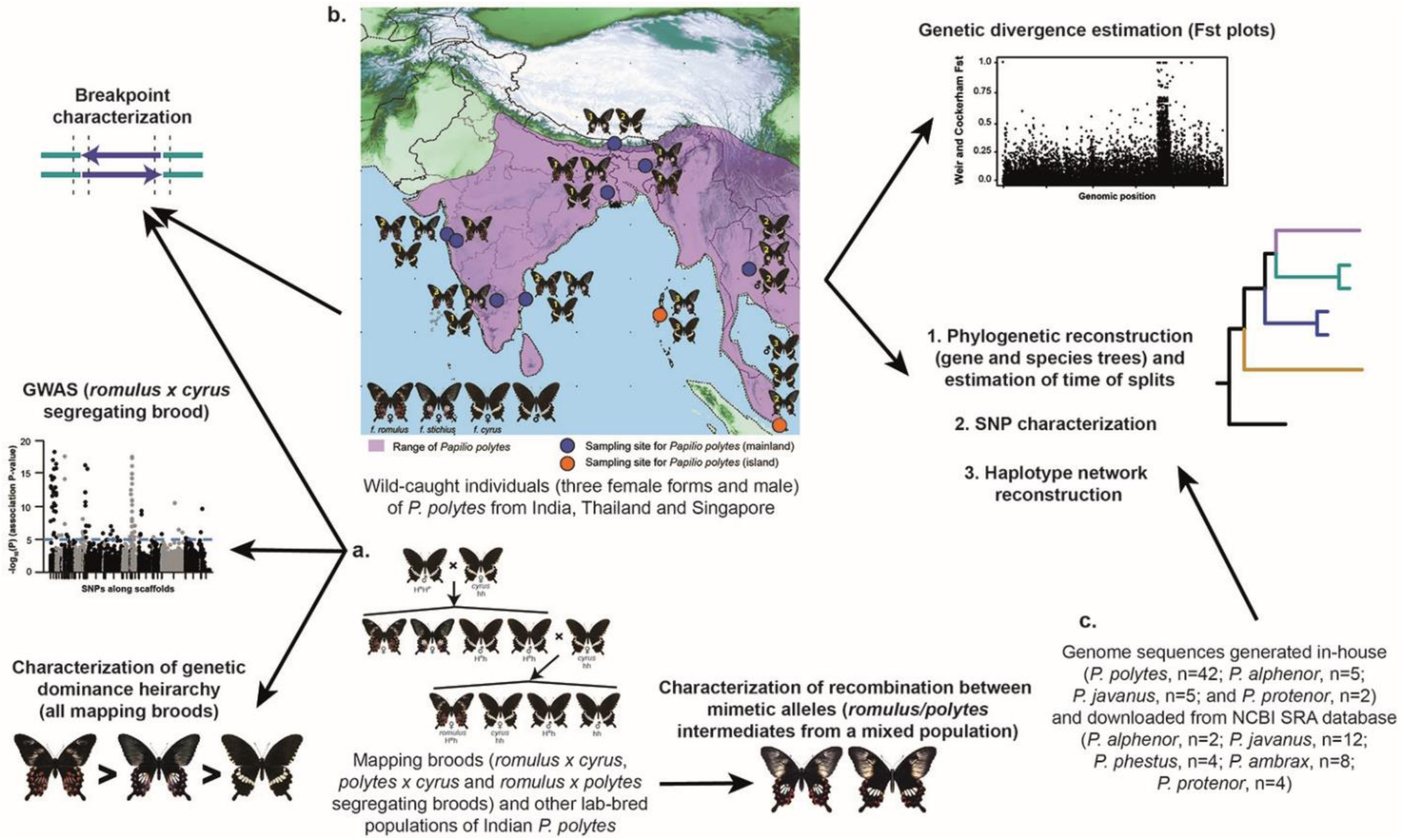
Sources of sequence data for the analyses performed in this study. *Papilio polytes* distribution, as determined from the tree topology in Fig. 1, is shaded in purple (b). Numbers on butterfly images indicate sample sizes for genome sequences of each female form/male from the respective sampling site.

**Figure S2:**
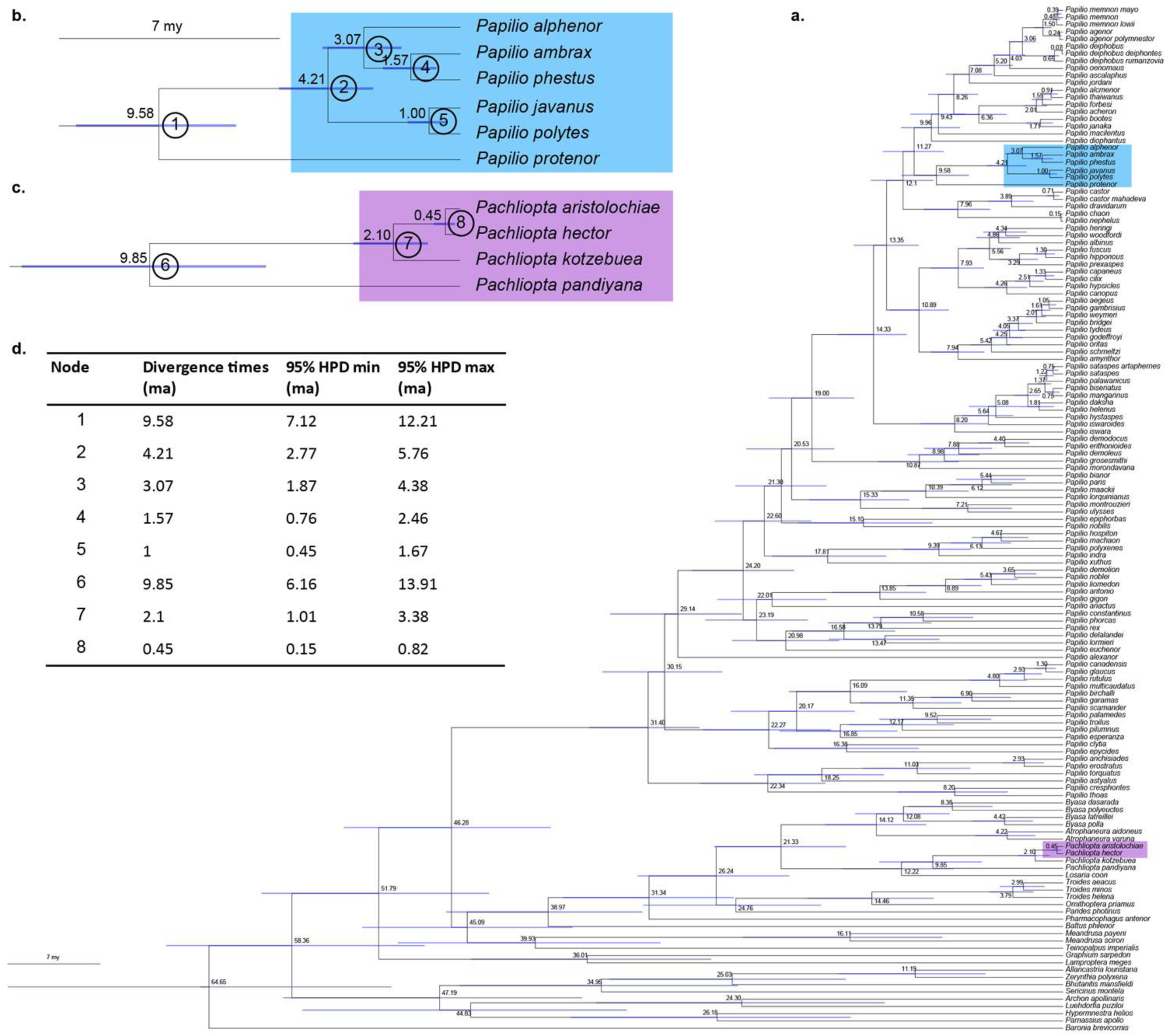
Time calibrated phylogeny and divergence times of models and mimics. **a.** The *Papilio* phylogeny showing node ages for all ingroups and outgroups considered in this work. The *Papilio polytes* species group and its *Pachliopta* aposematic models are highlighted in blue and purple, respectively. Numbers at nodes represent divergence times. All divergence times are in million years. **b.** The *P. polytes* species group, and **c.** the aposematic *Pachliopta*, are enlarged from panel **a**, with node ages represented on the top left corner of the nodes, and node numbers used in panel ‘d’ circled on the right side of the node. **d.** Node numbers, corresponding divergence times and 95% HPD min-max ranges are shown for all *Papilio polytes* species group species and their *Pachliopta* models to compare their relative ages.

**Figure S3:**
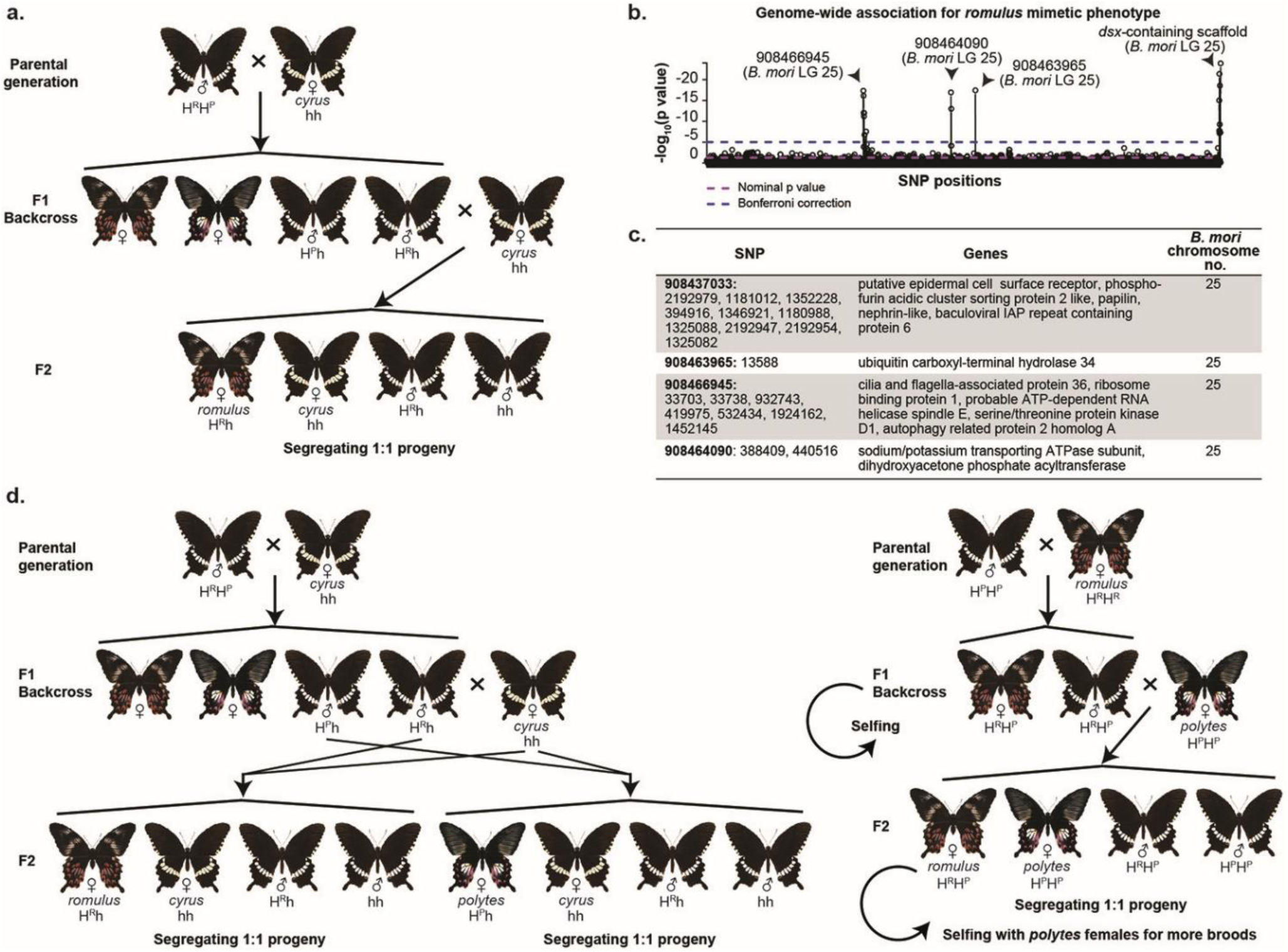
Genotype-phenotype association between *dsx* and *f. romulus.* a. The cross designed to generate a *romulus-cyrus* segregating brood to map the mimetic locus associated with *f. romulus*. b. Loci significantly associated with *f. romulus* and the scaffolds they lie on in the *P. polytes* genome (n=102: two parents and 50 females each of *f. cyrus* and *f. romulus*). The dotted lines indicate significance thresholds of p-values. c. The positions of all loci significantly associated with the mimetic phenotype and the genes in which they lie. Chromosome 25 of *Bombyx mori* contains *dsx*. d. Cross design to clarify dominance hierarchy (sample sizes given in Extended Data Table 2).

**Figure S4:**
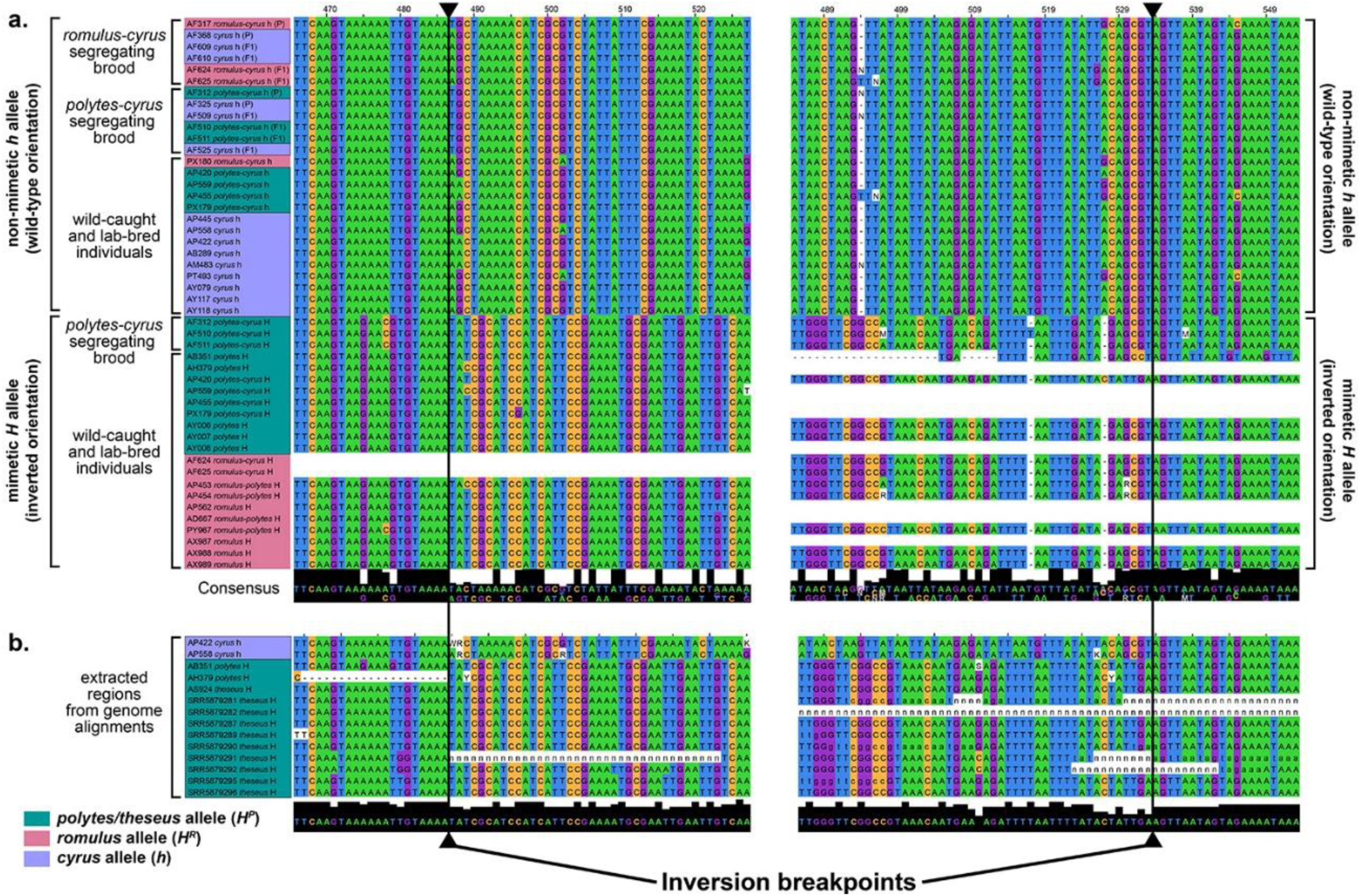
Inversion breakpoints for *H^R^* and *H^P^* alleles in *P. polytes.* a. DNA sequence alignment of the *dsx* inversion breakpoints from Sanger sequencing data, where blank spaces in rows indicate that PCRs did not work for the specific samples or breakpoints. The left and right blocks of sequences represent the left and right breakpoints of the *dsx* inversion. The details of each sequence are given on the left, and colour coded by female forms. The black row at the bottom includes consensus sequence, with the most prevalent base depicted at the top. Height of this row indicates sequence similarity. Inside the breakpoints, sequence similarity degrades, and the height of the black row at the bottom decreases. The precise breakpoints are marked with solid black lines and arrowheads. Inversion breakpoints for both the mimetic forms are identical. b. Genomic alignments showing *dsx* inversion breakpoints in *f. theseus*.

**Figure S5:**
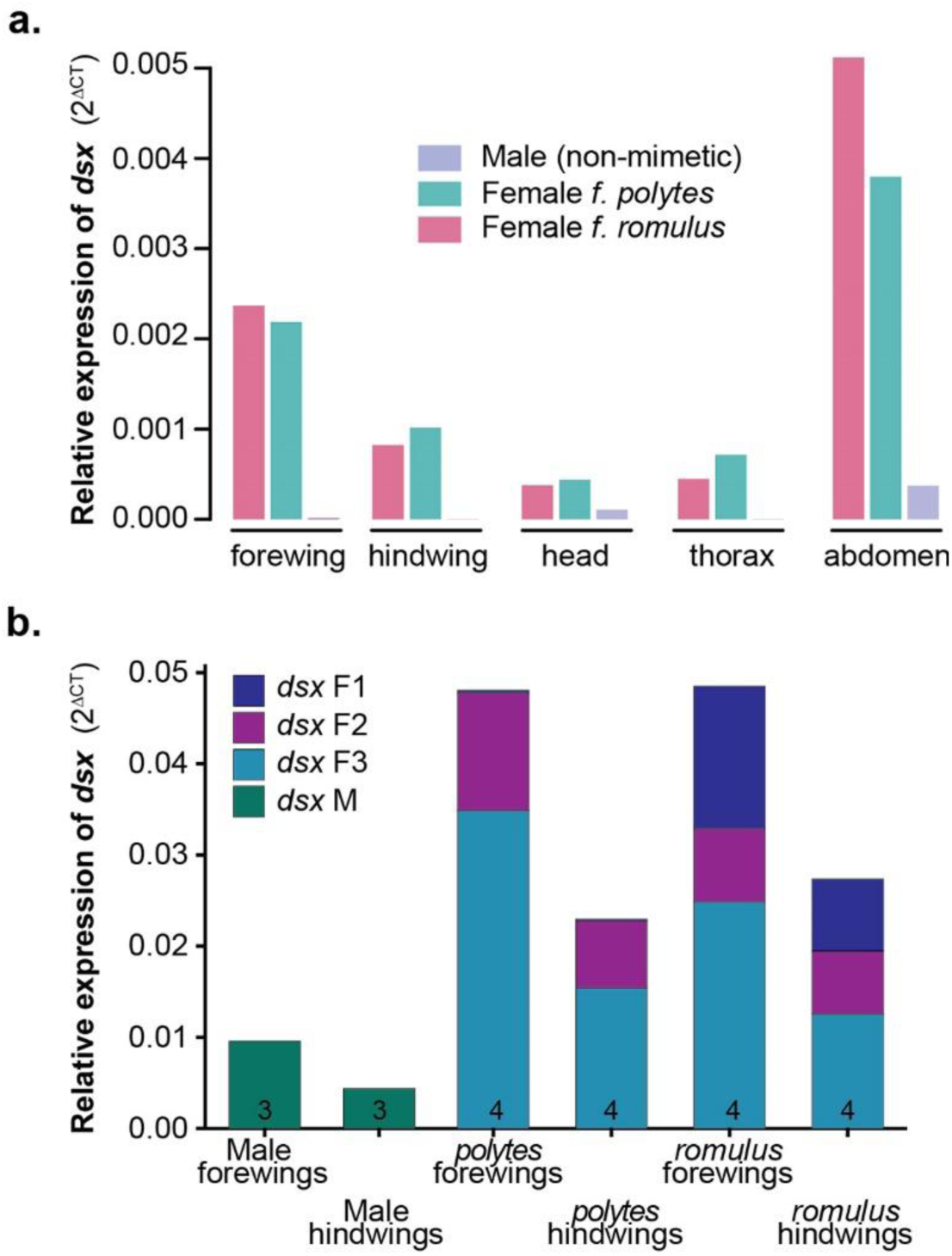
Expression of *dsx* isoforms in developing mimetic and non-mimetic wing patterns. Sex- and tissue-specific expression of *dsx* gene (panel a) and isoforms (panel b) across female forms (mimetic wings) and males (non-mimetic wings) in 3-day old pupae. Numbers in panel b indicate sample sizes for each female form and tissue. The composition of sex-specific *dsx* isoforms is shown in Extended Data Figure 6.

**Figure S6:**
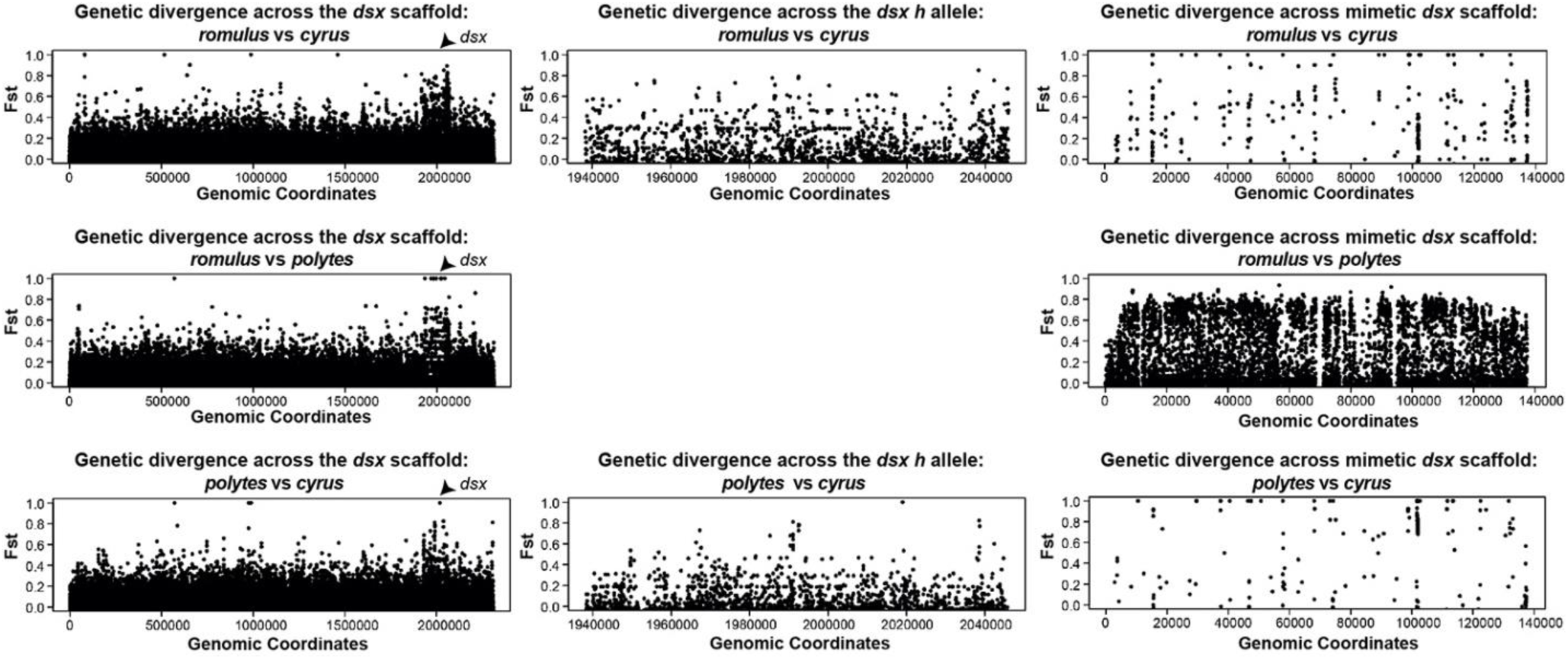
**Genetic divergence in *dsx-*containing scaffolds of the three *P. polytes* female forms.** From left to right, the figure shows plots of genetic divergence (Fst) between the non-mimetic *dsx* scaffold, the non-mimetic *dsx* gene and the mimetic *dsx* scaffold. The left panels highlight the *dsx* gene as a high Fst peak across the entire non-mimetic scaffold. Middle panels zoom into this high Fst peak to show several sites of high divergence in the pairwise comparisons of the three female forms. The right panels depict the mimetic scaffold that only contains *dsxH*, and shows high Fst peaks in the *romulus-polytes* comparison. Sample sizes for the three *dsx* alleles: *H^R^*=10; *H^P^*=13; and *h*=13.

**Figure S7:**
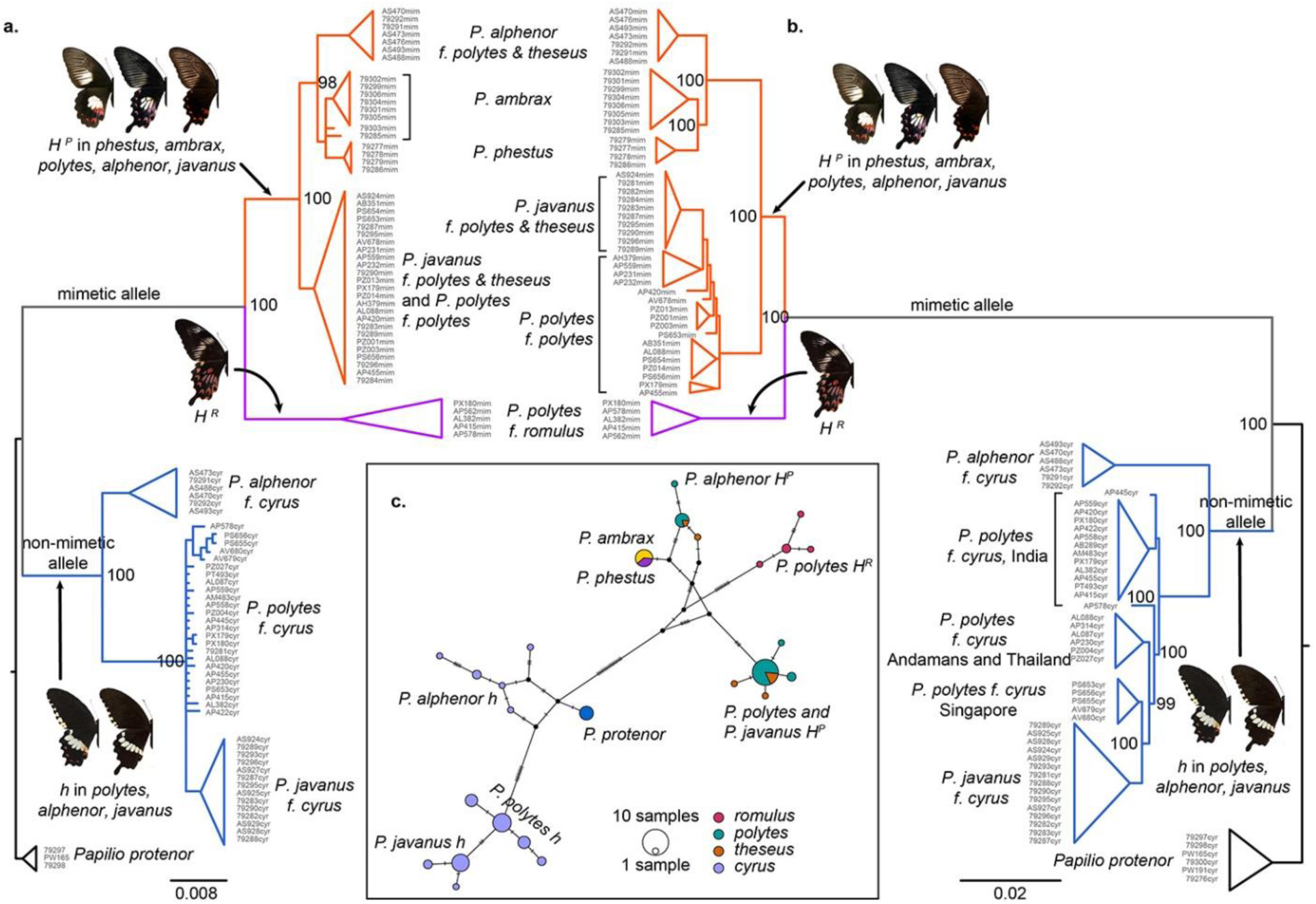
Evolution of *dsx* alleles in the *polytes* species group. We reconstructed gene trees using the CDS (a) and the non-coding regions of *dsx* (b), with *P. protenor* as the outgroup. The *dsx* alleles are colour coded, and the mimetic and non-mimetic female forms they produce are depicted (see Table S1 and Fig. 1 for *dsx* allelic abbreviations and female forms). c. Haplotype network generated with complete *dsx* CDS. Each female form clusters across species in both gene trees and haplotype network, indicating that the allelic polymorphism of *dsx* is much older than the individual species in which the alleles occur.

**Figure S8:**
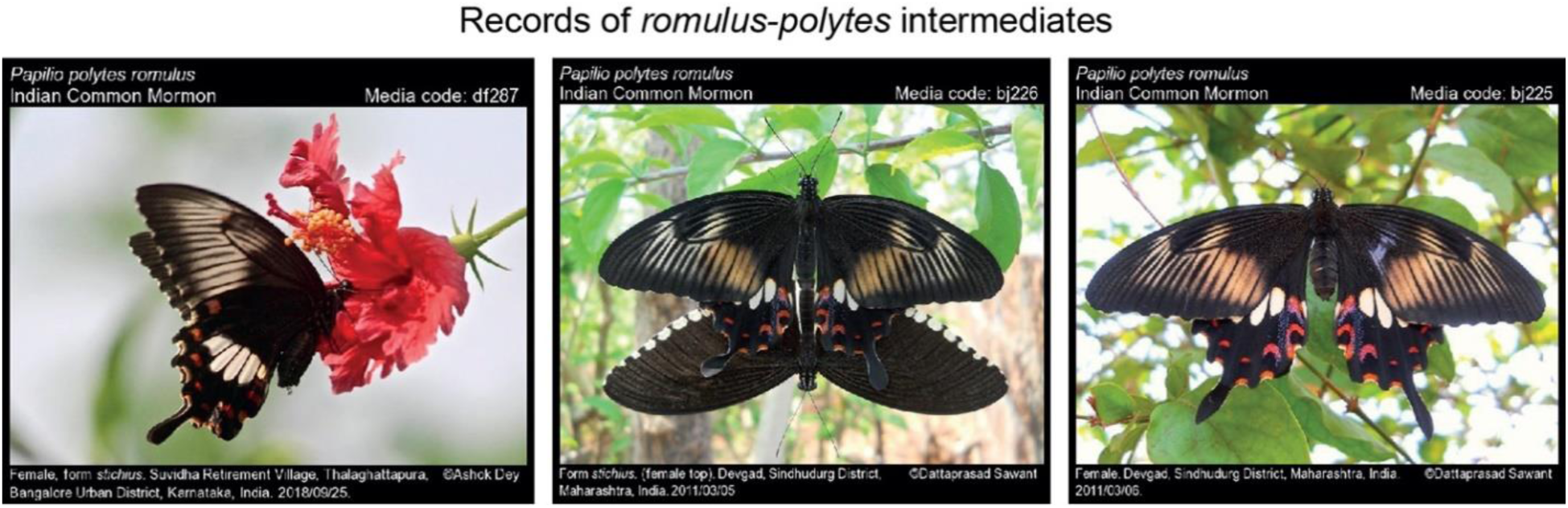
Occurrence of *romulus-polytes* intermediates in nature. The phenotypic intermediates possessing *romulus*-like forewings and *polytes*-like hindwings occur in nature, two of which were genetically characterized in Fig. 5. Image courtesy: Ashok Dey and Dattaprasad Sawant, from the Butterflies of India citizen science project (http://www.ifoundbutterflies.org/sp/603/Papilio-polytes).

